# Trophoblast and blood vessel organoid cultures recapitulate the role of WNT2B in promoting intravillous vascularization in human intrauterine and ectopic pregnancy

**DOI:** 10.1101/2022.04.18.488605

**Authors:** Xiaoya Zhao, Zhenwu Zhang, Yurui Luo, Qinying Ye, Shuxiang Shi, Xueyang He, Jing Zhu, Qian Zhu, Duo Zhang, Wei Xia, Yiqin Zhang, Linlin Jiang, Long Cui, Yinghui Ye, Yangfei Xiang, Junhao Hu, Jian Zhang, Chao-Po Lin

**Affiliations:** Department of Obstetrics and Gynecology, International Peace Maternity and Child Health Hospital, School of Medicine, Shanghai Jiaotong University, No. 910, Hengshan Rd, Shanghai, 200030, China; Shanghai Municipal Key Clinical Specialty, Shanghai, China; School of Life Science and Technology, ShanghaiTech University, Shanghai, 201210, China; Interdisciplinary Research Center on Biology and Chemistry, Shanghai Institute of Organic Chemistry, Chinese Academy of Sciences, Shanghai, 201210, China; Department of Reproductive Endocrinology, Women’s Hospital, Zhejiang University School of Medicine, Hangzhou, 310000, China.

**Keywords:** Trophoblast organoid, blood vessel organoid, angiogenesis, ectopic pregnancy, WNT signaling

## Abstract

Tubal ectopic pregnancy (TEP), a pregnancy complication caused by aberrant implantation in fallopian tubes, accounts for 9-13% pregnancy-related deaths. The lack of models for human TEP hampers the understanding of its pathological mechanisms. Here, we employed multiple models to investigate the crosstalk between human trophoblast development and intravillous vascularization. We found that the severity of TEP, the size of placental villi, and the depth of trophoblast invasion are correlated with the extent of intravillous vascularization. We identified a key pro-angiogenic factor secreted by trophoblasts, WNT2B, that promotes villous vasculogenesis, angiogenesis, and vascular network expansion. In an organoid coculture model consisting of trophoblast organoids and blood vessel organoids, knockdown of WNT2B in trophoblast organoids compromises their pro-angiogenic effect on the development of blood vessel organoids. These organoid-based models reveal an important role for WNT-mediated angiogenesis in pregnancies and could be employed to investigate the commutations between trophoblasts and endothelial/endothelial progenitor cells.

## Introduction

Ectopic pregnancies (EP) occur when embryos implant at sites other than uterine endometria (intrauterine pregnancies, IP), with 98% of EP located in the fallopian tubes, while others can occur in the ovaries or abdominal cavities (Marion and Meeks, 2012). The incidence of EP is approximately 2% of all reported pregnancies, although the actual rate could be higher due to misdiagnoses (Bronson, 2018; Farquhar, 2005). Despite substantial improvements in diagnosis and management, EP remains a significant cause of maternal morbidity and mortality in early pregnancies, accounting for 9-13% of all pregnancy-related deaths (Farquhar, 2005). Tubal EPs (TEP) can be further classified into two major types: abortive or ruptured ectopic pregnancies (hereafter AEP and REP, respectively) (Caspi and Sherman, 1987). These two forms can be diagnosed or confirmed by human chorionic gonadotropin (hCG) levels, transabdominal ultrasonography, and laparoscopy (Alkatout et al., 2013; Doubilet et al., 2014). AEPs are frequently characterized by low and continuously decreasing hCG levels and hemodynamic stability of unruptured fallopian tubes (2013). In contrast, REPs are accompanied with high and continuously increasing hCG levels (>3000-5000 IU/L), active fetal hearts, and maternal hemodynamic instability. Several risk factors of EPs have been reported (Farquhar, 2005; Fukami et al., 2016; Hendriks et al., 2020; Marion and Meeks, 2012). However, whether these factors share a causal relationship with REP has not yet been substantiated, reflecting a lack of understanding regarding the mechanism(s) underlying the distinct pathologies of AEP and REP.

Analogous to the extensive trophoblast invasion into fallopian tube walls in REP, trophoblasts invade deeply into uterine decidua in IP during the first trimester, reaching as far as the myometrium, and remodel the decidual vasculature to increase the maternal blood supply (Hemberger et al., 2020; Sato, 2020). Invading trophoblast cells differentiate into the outer layer syncytiotrophoblasts (STBs) and inner layer villous cytotrophoblasts (vCTBs, the stem cell population in trophoblasts), thus forming primary villi (Sato, 2020). After day 15 post conception (p.c.), extraembryonic mesoderm cells migrate into primary villi to form the stromal centers of secondary villi (Sato, 2020). Between day 18 and 20 p.c., those mesenchymal cells differentiate into endothelial cells (ECs), forming fetal capillaries, which are hallmarks of tertiary villi development (Sato, 2020). The developed placental villi form the fetal side of the maternal-fetal interface which functions as a barrier between maternal and fetal blood while also allowing the exchange of gas, nutrients, and waste between the mother and the fetus (Hemberger et al., 2020).

Vasculogenesis, angiogenesis, and vascular network development in placental villi are crucial for establishing a viable pregnancy (Kaufmann et al., 2004; Wang and Zhao, 2010). Villous vasculogenesis refers to the differentiation of multipotent mesenchymal cells into hemangioblastic progenitor cells, which ultimately form *de novo* primitive capillary networks (Kaufmann et al., 2004; Wang and Zhao, 2010). In the following stage, villous angiogenesis, placental capillary networks form via sprouting and elongation of preexisting blood vessels in villous trees (Kaufmann et al., 2004). Considering the importance of circulation in the functional maternal-fetal interface, it is unsurprising that abnormalities in the placental vascular bed underpin a variety of pregnancy complications, including placental chorangiosis, spontaneous miscarriage, preeclampsia, gestation diabetes, and intrauterine growth restriction (Barut et al., 2012; Santa et al., 2015; Soma et al., 2013; Verlohren and Dröge, 2020). Aberrant expression of several pro-angiogenic factors, such as VEGF (vascular endothelial growth factor), PlGF (placental growth factor), sFlt-1 (soluble fms-like tyrosine kinase 1), fibroblast growth factors (FGF), and angiopoietins (ANG), is associated with placental vascular dysfunction and pregnancy complications (Plaisier et al., 2007; Sherer and Abulafia, 2001). Hence, abnormalities in vasculogenesis and angiogenesis can impact the function of trophoblasts, the depth of trophoblastic invasion, as well as the growth of the embryonic and extraembryonic fetal tissues.

Investigations on the communications between trophoblasts and endothelial cells/endothelial progenitor cells in human are largely hampered by the lack of proper cell culture models and by the differences between species (*i.e.*, structural differences of placentas between rodents and primates, as well as the absence/rarity of EP phenotypes in rodents (Benirschke et al., 2012; Corpa, 2006; Furukawa et al., 2014)). In this study, by combining multiple vasculogenesis/angiogenesis analyses and blood vessel/trophoblast organoid cultures, we found that secreted factors from placental villi or trophoblast organoids promote vasculogenesis and angiogenesis in IP and REP conditions. This pro-vasculogenic/angiogenic effect is likely mediated by trophoblast-secreted WNT2B which promotes the differentiation of endothelial cells, activates VEGF, and vascular network development to support fetal growth. Collectively, our models and results identify the WNT signaling as a mediator of vasculogenesis, angiogenesis, and network formation during early pregnancy, thus shedding light on intricate interactions between trophoblasts and villous endothelial cells/endothelial progenitor cells, suggesting its use a diagnostic marker and a potential therapeutic target for TEP interventions.

## Results

### Morphological and immunological similarity between TEP and IP placental villi

In IP and TEP conditions, embryos implant and develop in the decidualized uterine endometrium and the fallopian tube wall, respectively. Since only the decidualized endometrium has adequate physical space, blood supply, and the appropriate microenvironment to support optimal fetal growth (Liu et al., 2020; Ng et al., 2020), it is unclear why implanted embryos still exhibit extensive trophoblast invasion and continuous fetal growth that together lead to tubal rupture in ectopic pregnancy (REP). We therefore hypothesized that the maternal-fetal interface associated with REP could share more similarities with IP than with tubal abortive ectopic pregnancy (AEP). To explore this possibility, we selectively collected IP, AEP, and REP placental villi for further analyses (Figure 1A). All clinical samples were devoid of medical treatments or obvious risk factors associated with fallopian tubal abnormalities, *e.g.*, smoking, obvious tubal inflammatory adhesions, previous fallopian tubal diseases, and history of tubal surgery (Farquhar, 2005; Marion and Meeks, 2012). No significant differences were found between the three groups in terms of maternal age, gravity, parity, or gestational days (Figure S1A).

**Figure 1.**
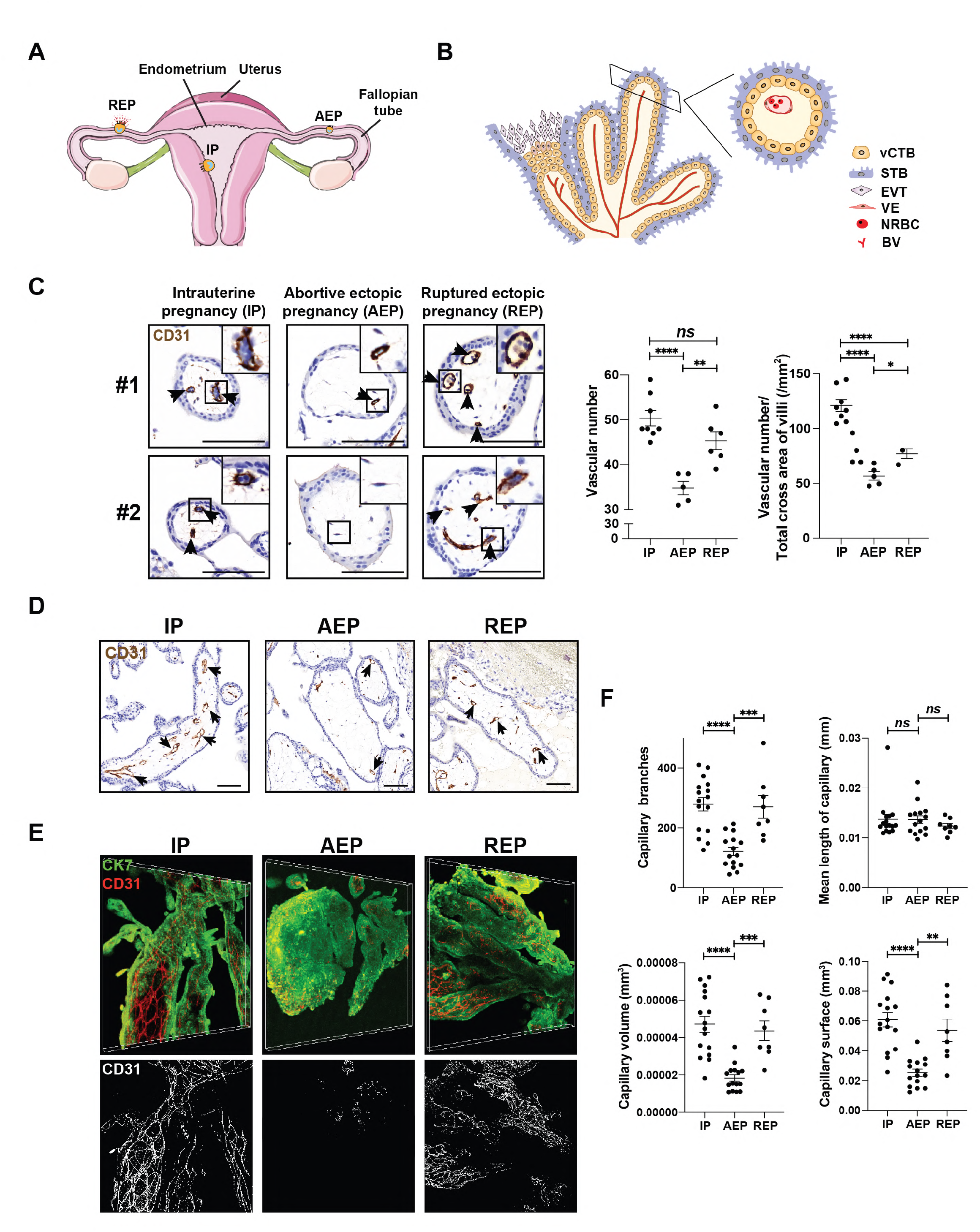
IP, AEP and REP placental villi exhibit major differences in intravillous vascularization. (A) Schematic illustration (left) and representative macroscopic images (right) of three types of pregnancies. Left, in intrauterine pregnancy (IP), embryos are implanted on the endometrium of the uterus. In both abortive pregnancy (AEP) and ruptured pregnancy (REP), embryos are implanted on fallopian tubes, yet only REP exhibits excessive trophoblast invasion and overgrowth of fetuses, leading to the rupture of tubal walls. (B) Schematic illustration images of the placental villus. In the first trimester of pregnancy, the trophoblastic column of placental villus is consisted of intervillous capillaries and stromal cells, both of which are surrounded by trophoblast shells made of syncytiotrophoblasts (STBs) and villus cytotrophoblasts (vCTB). In anchoring villi, vCTBs protrude the STB layer and differentiate into extravillous trophoblasts (EVTs). VE: vascular endothelium, NRBC: nucleated red blood cell, BV: blood vessel. (C) Left, representative images of immunohistochemical staining for CD31 to identify intravillous capillaries (arrows) in cross sections of IP (n=8), AEP (n=5), and REP (n=6) placental villi. Right, quantification of vascular numbers and the ratio of vascular numbers to total cross area of villi in three types of pregnancies. Each dot represents a clinical sample. AEP placental villi contain much less capillaries and fewer branches per unit area than REP, which present no significant difference in vascular numbers compared with IP. Scale bars, 100 μm. Data represents mean ± SEM. (D) Immunohistochemical staining of CD31 in longitudinal sections of IP, AEP, and REP placental villi. Scale bars, 100 μm. (E-F) IP and REP placental villi exhibit higher vascularization level in cleared tissues. 16 samples of IP, 15 samples of AEP, and 8 samples of REP placental villi were cleared and immunostained for CK7 (green, trophoblasts) and CD31 (red, endothelial cells). Stereoscopic views of one representative sample for each type of pregnancy are shown (see Figure S2 for the additional representative sample). Lower, skeletonization of CD31 staining to visualize vascular systems of IP, AEP and REP placental villi with quantification of capillaries in cleared placental villi from three types of pregnancies (F). Each dot represents a clinical sample. AEP (n=15) placental villi contained compromised vascular systems compared with IP (n=16) in the first trimester of pregnancy in number of branches, diameters, and volumes, while REP (n=8) placental villi possessed vascular structures more resembling IP condition. Scale bars, 100 μm. Quantification results are represented as mean ± SEM. \**P* < .05, \*\**P* < .01, \*\*\**P* < .001, \*\*\*\**P* < .0001. *P* values were calculated by ANOVA, Tukey’s test.

Among the three pregnancy types, hCG levels in the maternal circulating blood of the IP group was ∼5 fold higher (47949.79±6412.00 IU/L) than that of the REP group, while REP patients exhibited ∼10 fold higher levels (7178.78±482.50 IU/L) than that detected in AEP patients (858.83±64.77 IU/L) (Figure S1A), in agreement with previous reports that hCG levels are correlated with tubal rupture (Goksedef et al., 2011). We also compared the volumes of individual villous trees among the three pregnancy groups and found that the total volume was obviously greater in the IP group, with more defined dendritic structures than those from AEP and REP patients (Figure S1B). The total volume of individual villous trees from REP was 2-3 times larger than that of AEP patients (Figure S1B). Notably, same trend was observed in fetus size among groups (Figure S1C). In addition, heart activity was detected in 20/28 REP fetuses, whereas 0/34 AEP fetuses had active hearts, indicating fetal development was more advanced in REP (Figure S1A). Taken together, these observations suggested that at similar gestational ages, REP was accompanied by more extensive embryonic and extraembryonic tissue development than AEP.

### Major differences in vascular systems among IP, AEP, and REP placental villi

In normal IP, functional vasculature develops in the placental villi in four stages: primary (11-13 days p.c.), secondary (16^th^ day-3^rd^ wk p.c.), tertiary (3^rd^ wk-4^th^ month p.c.), and free villi (> 4^th^ month p.c.). At the tertiary stage, the mesenchymal tissue begins to differentiate into endothelial cells (ECs) to form capillaries which, together with trophoblasts, constitute the hemochorial placental barrier (Blundell et al., 2016; Sato, 2020) (Figure 1B). We found that tertiary placental villi from the IP and REP groups appeared to have more capillary-like tubules than those of the AEP group as demonstrated by immunohistochemical staining for CD31 (PECAM) on cross and longitudinal sections of IP, AEP, and REP villi to compare development of the EC layer (Figure 1C-D, and Figure S2A for additional samples). AEP placental villi also contained significantly fewer and smaller capillaries per unit area than REP villi, which shared greater similarity in vascular characteristics with IP (Figure 1C).

To exclude the possibility that CD31 labelled lymphatic tissues rather than blood vessels, we also used an immunohistochemical stain for the lymphatic endothelium marker D2-40 (podoplanin) in IP and TEP villi. We detected no D2-40 signal in placental villi of either IP or TEP, while a strong D2-40 signal was evident in cervical lymph node positive control samples (Figure S2B), which indicates that CD31-positive cells in placental villi were exclusively vascular, not lymphatic, ECs.

Since analysis of 2D sections of villous trees can be limited by the integrity of clinical samples and section regions, we performed tissue clearing on all three types of placental villi, stained whole-mounts for CK7 and CD31 (Merz et al., 2018), and compared villi volumes, branching patterns and morphology of blood vessels. Three dimensional analyses of cleared tissues confirmed that AEP placental villi exhibited compromised vascular systems, *e.g.*, lower number of branches, vessel diameter, and volume of blood vessels, compared with IP villi collected at the same gestational stage (Figure 1E-F, Figure S2C for additional samples, and Video S1-6). In line with our findings above, the vascular structure of REP placental villi more closely resembled those of IP patients (Figure 1F). The highly developed vascular systems observed in both IP and REP placental villi suggested greater transport and availability of oxygen and nutrients from the ectopic microenvironment to support fetal development.

### Transcriptomic differences in signal pathways and developmental markers between IP and TEP placental villi

To identify transcriptomic differences between IP and TEP placental villi that underlie ectopic, rather than eutopic, pregnancy, we conducted RNA-seq analysis for five placental villi each from IP and TEP patients, using the same inclusion criteria as above. Principal component analysis (PCA) and unsupervised clustering showed that samples clustered into distinct IP and TEP groups, which suggested substantial differences in gene expression between placental villi in intrauterine and tubal pregnancies (Figure 2A-B).

**Figure 2.**
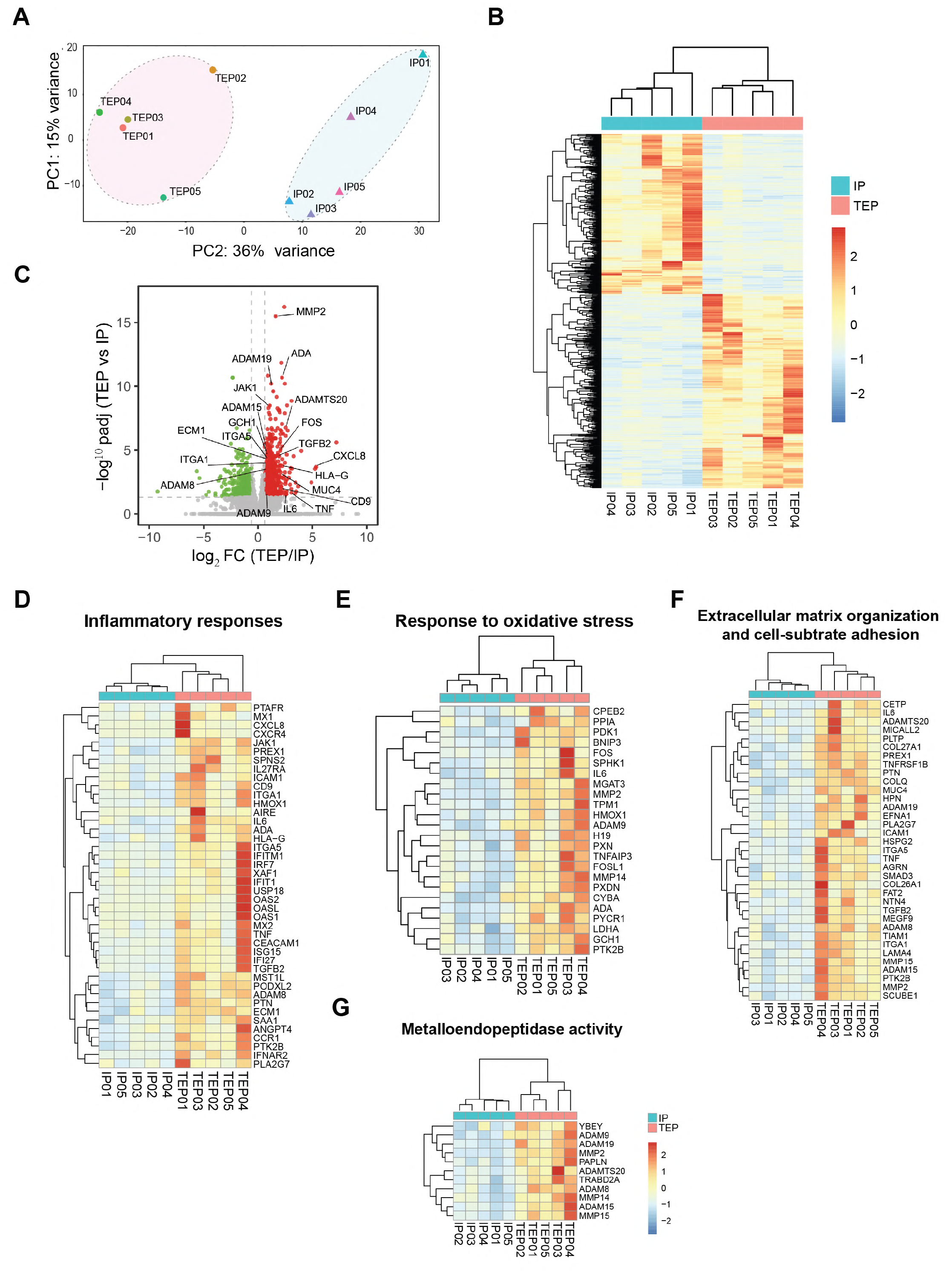
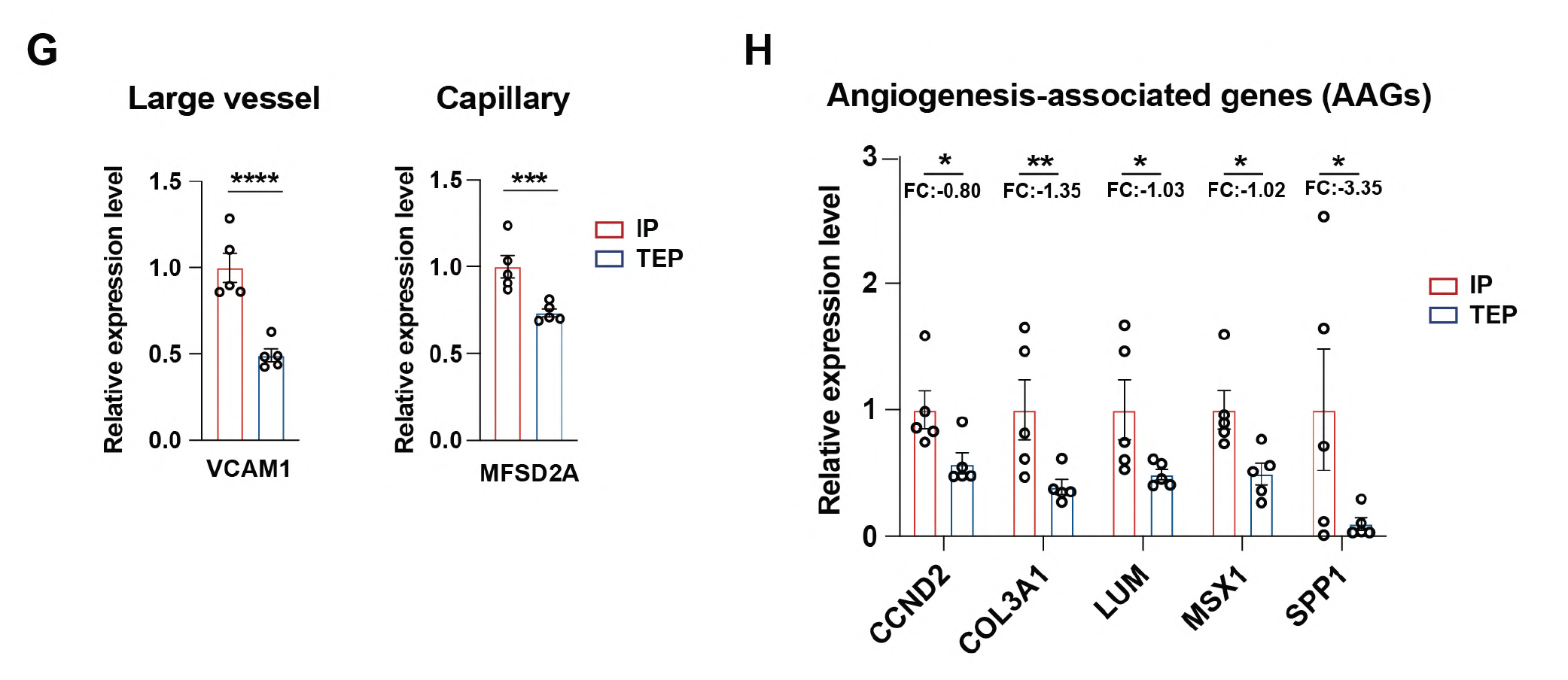
Transcriptome analyses of signaling pathways and developmental states in IP and TEP conditions. (A, B) Transcriptomes of placental villi from intrauterine pregnancy (IP, n=5) and tubal ectopic pregnancy (TEP, n=5) are segregated into two groups, as demonstrated by principal component analysis (PCA) (A) and unsupervised hierarchical clustering based on top 1,123 differentially expressed genes (DEGs) (B). Oval represents confidence intervals for IP (triangle) and TEP (circle). (C) The volcano plot for comparison of IP (n=5) and TEP (n=5) transcriptomes. Red dots represent up-regulated genes, green dots represent down-regulated genes, and the grey dots represent genes with no significant differences in expression levels. Important DEGs revealed in pathway analyses are annotated. (D-G) Pathway-specific enrichment of genes up-regulated in TEP. (F-G) Expression of markers of large vessels and capillaries (F) and angiogenesis-associated genes (G) are down-regulated in TEP samples. \*\**P* < .01, \*\*\**P* < .001, \*\*\*\**P* < .0001. *P* values were calculated by student’s *t*-test or Welch’s *t*-test.

We then performed GO and GSEA analyses on the differentially expressed genes (|log_2_ FoldChange| ≥ 0.6 and *P* value ≤ 0.05) to determine which signaling pathways and/or developmental programs were enriched in TEP (Figure 2C and S3A). The analysis suggested that expression of inflammation-related genes was elevated in TEP samples, consistent with a proposed link between inflammation and tubal implantation (Figure 2D and S3B) (Dekel et al., 2014; Granot et al., 2012). We also found that hypoxia-inducible genes in the oxidative stress-related gene set were up-regulated in the TEP villi samples (Figure 2E and S3C), possibly due to the elevated inflammatory response or compromised angiogenesis related to low hCG (Lam et al., 2004; Roth et al., 2010). In addition, extracellular matrix (ECM) organization- and cell adhesion-related genes were enriched in TEP placental villi (Figure 2F and S3D) which, together with upregulation of metalloendopeptidases (Figure 2G and S3E), suggested dysregulation of the interactions between trophoblasts and the microenvironment (tubal walls). Moreover, the expression of markers for large vessels and capillaries was decreased in TEP villi (Figure 2H), which was in agreement with the inhibited vascularization phenotype of TEP placental villi (Figure 1C-F and S2). Besides of blood vessel markers, angiogenesis-associated genes (AAGs) (Qing et al., 2022) were also down-regulated in TEP samples which, again, suggesting the compromised angiogenesis process (Figure 2I). These results showed substantial differences in gene expression patterns between placental villi that developed in uteri versus fallopian tubes, despite their similar morphologies and trophoblast organizations (see later).

### Secretory factors from placental villi promote angiogenesis

To next investigate whether and which angiogenic factors may contribute to aberrant vascularization in TEP villi, we performed 2D tube formation and 3D sprouting assays to visualize luminal, capillary-like development using human umbilical vein endothelium cells (HUVECs) (Ko and Lung, 2012; Zahra et al., 2019). Briefly, we conducted 2D and 3D angiogenesis assays by culturing HUVECs as monolayers (tube formation assay) or spheroids (sprouting assay) in media conditioned with either fetal or maternal explants from each of the three pregnancy types (Figure 3A). Treatment with VEGF strongly induced the formation of tube networks or vascular sprouts in both 2D and 3D assays, respectively (Figure 3B and 3C), which indicated that these platforms are robust for examining angiogenesis.

**Figure 3.**
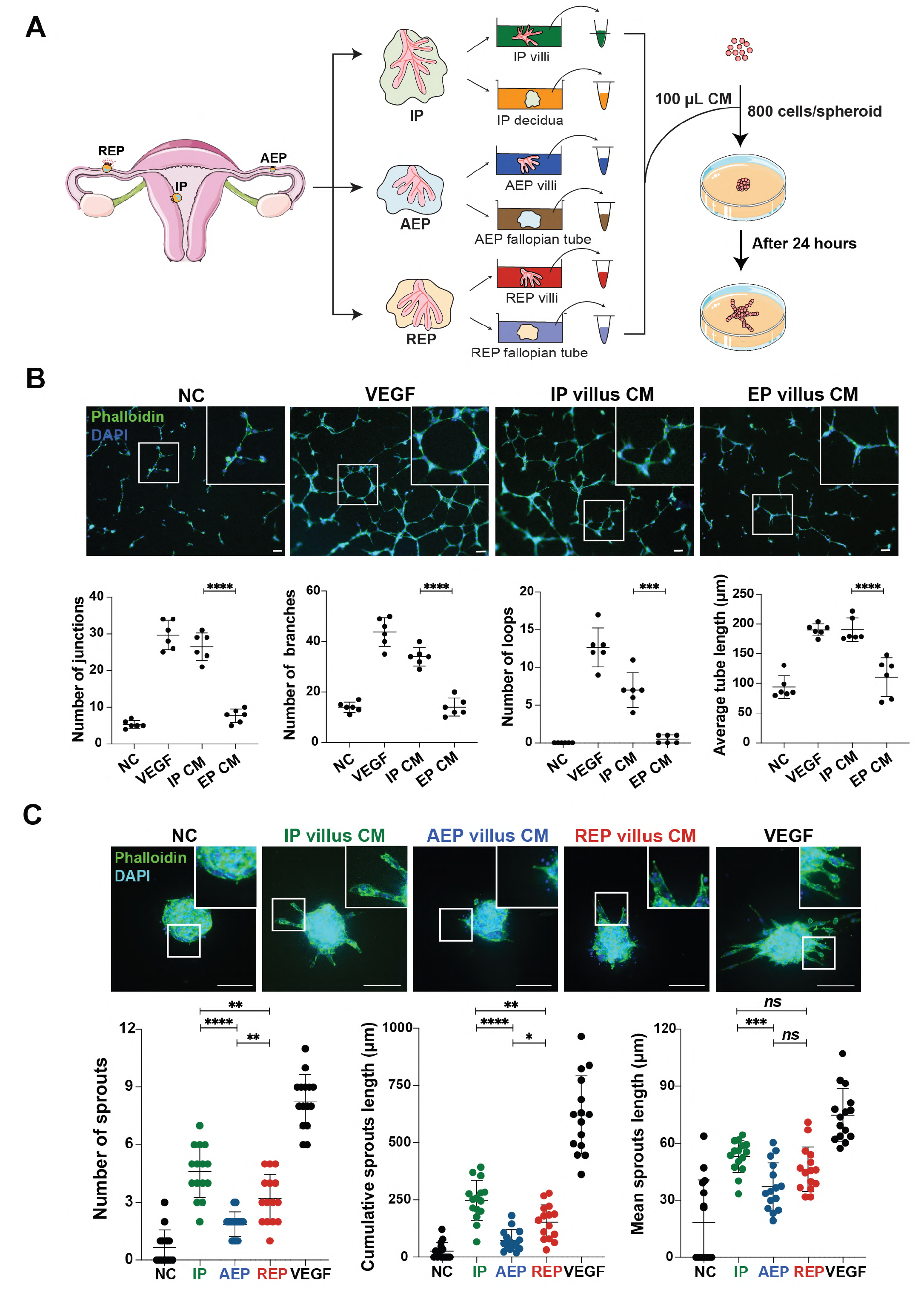
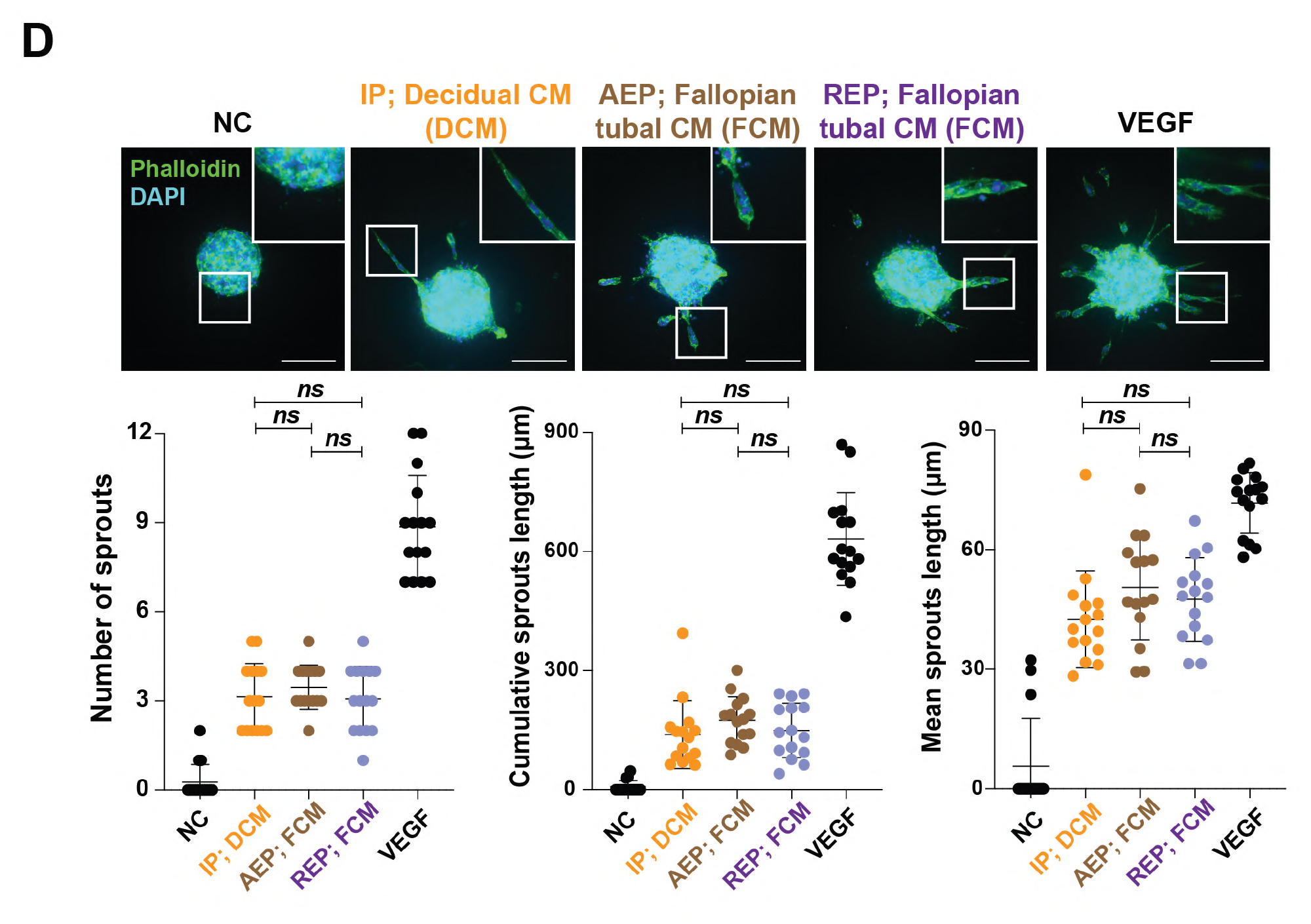
Secretory factors from placental villi promote angiogenesis. (A) Schematic workflow for sprouting angiogenesis assay. IP: intrauterine pregnancy (IP), AEP: abortive pregnancy, REP: ruptured pregnancy, CM: conditioned medium. (B-D) Secretory factors of the fetal part, not maternal part, from the maternal-fetal interface promotes sprouting of blood vessels *in vitro*. (B) CM from IP, but not EP, promotes the tubal formation of HUVECs. (C-D) Conditioned media were collected from fetal (placental villi) (C) or maternal (endometrium in the IP condition and fallopian tube wall in the AEP/REP condition) (D) and explants and tested for their angiogenic potential by sprouting assays. VEGF (250 ng/ml) serves as the positive control. Phalloidin/DAPI staining was employed to enhance the contrast. Representative images and the quantitative data of the number of sprouts per spheroid, as well as the mean/cumulative sprout length are shown. Each dot represents a field of view and values from fifteen independent fields of view are represented as mean ± SD Scale bars, 100 μm. \**P* < .05, \*\**P* < .01, \*\*\**P* < .001, \*\*\*\**P* < .0001. *P* values were calculated by ANOVA, Tukey’s test and student’s *t*-test or Welch’s *t*-test. NC: negative control, DCM: decidua conditioned medium, FCM: fallopian tubal conditioned medium.

To determine whether angiogenic factor(s) are secreted from the maternal-fetal interface, as well as to determine their origin, we surgically isolated the maternal-fetal interfaces and separated their fetal (placental villi) and maternal (uterine endometria or fallopian tubal walls) components, cultured the explants, and harvested their respective conditioned media (Figure 3A). In all three types of pregnancies, conditioned media obtained from both fetal and maternal parts could promote angiogenesis in HUVEC-based tube formation and sprouting assays (Figure 3B and 3C). However, we only observed differences in pro-angiogenic potential between conditioned media of IP, AEP, and REP from fetal placental villi explants (Figure 3C), not from maternal tissues (Figure 3D). More specifically, compared to unconditioned medium, the conditioned medium from IP villi led to the strongest induction of tube network or sprout formation, followed by REP medium, then AEP medium, which was highly correlated with differences in placental villi vascularization observed by immunostaining (Figure 1C-E). These results led us to conclude that the factor(s) responsible for differences in vascular development within placental villi was secreted from the fetal compartment.

### IP and REP placental villi promote angiogenesis in a WNT-dependent manner

To identify which signaling pathway promotes angiogenesis in these placental villi, we re-examined the RNA-seq results and found multiple WNT pathway components that were downregulated in TEP placental villi (Figure 4A). Since some WNT ligands also exhibit pro-angiogenic activity in cancers, atherosclerosis, and eye diseases (Foulquier et al., 2018; Taciak et al., 2018; Wang et al., 2019), we hypothesized that WNT ligands secreted from placental villi could promote angiogenesis. To test this possibility, we used the β-catenin/CBP inhibitor ICG-001 (Akcora et al., 2018), as well as the tankyrase inhibitor XAV-939 (Jang et al., 2019), in combination with the conditioned media from IP, AEP, and REP, to interrogate the role of WNT signaling in angiogenesis using sprouting assay. Treatment with ICG-001 or XAV-939 significantly abolished the pro-angiogenic effects of conditioned media from IP or REP, suggesting that the angiogenic effects on HUVECs were canonical WNT signaling-dependent (Figure 4B and S4A). Consistent with this finding, treatment with other WNT antagonists, including secreted frizzled-related protein 1 (sFRP-1) and WNT inhibitory factor 1 (WIF-1) (Kawano and Kypta, 2003), also abolished the pro-angiogenic effects of IP- or TEP-conditioned media (Figure 4C and S4B). Notably, VEGF was almost undetectable in conditioned media from either IP or TEP placental villi, even at pg/ml ELISA sensitivity (Figure 4D). Since sFRP-1 and WIF-1 only inhibited pro-angiogenic effects of conditioned media (Figure 4C and S4B), not VEGF (Figure S5A), it’s less likely that VEGF-induced expression of WNT ligands in placental villus promoted angiogenesis. Taken together, these results suggested that the WNT signaling pathway directly promoted angiogenesis during pregnancy and could lead to vascular growth in REP condition.

**Figure 4.**
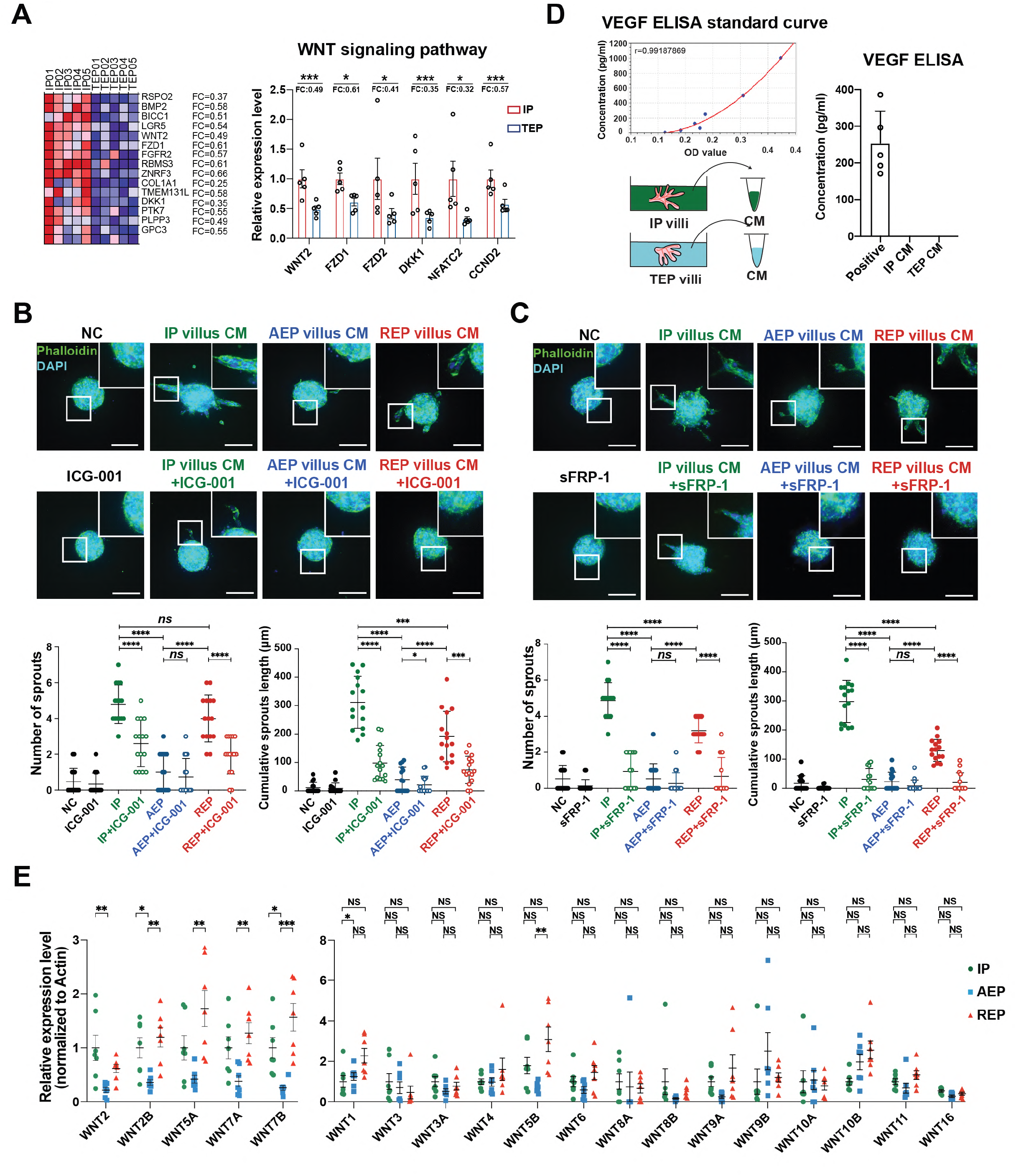
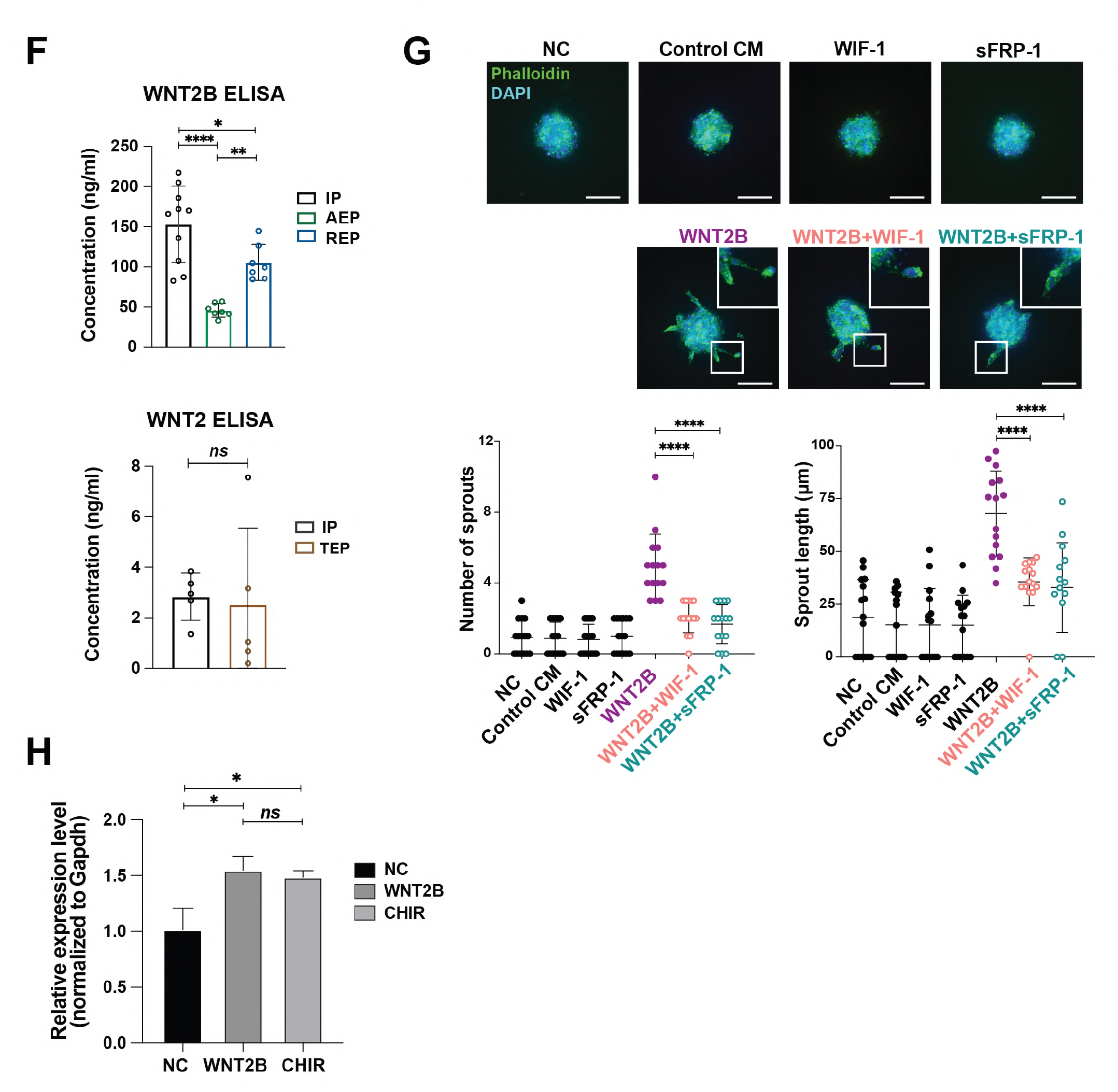
WNT2B activates VEGF and promotes angiogenesis. (A) Components of WNT signaling pathway are down-regulated in TEP (n=5) placental villi as compared with IP (n=5) ones. Expression levels of WNT-related genes, as revealed by GSEA analysis, are retrieved from RNA-seq data, log-transformed, normalized, and presented as the heatmap. Data represents mean ± SEM. *P* values were calculated by student’s *t*-test or Welch’s *t*-test. (B-C) Conditioned media from IP and REP placental villi explants promote angiogenesis in a WNT-dependent manner. In sprouting angiogenesis assay, conditioned media from three types of pregnancies were co-treated with varies inhibitors antagonizing different components of canonical WNT signaling pathway, such as ICG-001 (10 μM) (B) or sFRP-1 (5 μg/ml) (C). The number of sprouts per spheroid and the cumulative sprout length indicate that WNT inhibitors greatly abolish pro-angiogenic effects of conditioned media from IP and REP villous explants. (D) Absence of VEGF protein in conditioned media (CM) from IP (n=9) and TEP (n=9) placental villi as detected by ELISA assay. (E) Real-time qPCR analysis of all 19 human WNT ligands in IP (n=7) compared to AEP (n=7) and REP (n=7). Data represents mean ± SEM. (F) Upper, WNT2B protein level in the CM from the IP (n=10), AEP (n=7), and REP (n=7) placental villi, as detected by ELISA assay. Lower, WNT2 protein level in the CM from the IP (n=5) and TEP (n=5) placental villi, as detected by ELISA assay. (G) WNT2B’s pro-angiogenic activity is dependent on WNT signaling. In sprouting angiogenesis, capillary-like sprouts after 24 h stimulation of WNT2B conditioned medium with and without co-treatment of WIF-1 (2.5 μg/ml) or sFRP-1 (5 μg/ml) were imaged and quantified. The statistical data of number of sprouts per spheroid and cumulative sprout length indicates that sFRP-1 and WIF-1 significantly reduces the pro-angiogenic effects of WNT2B. (H) Treatment of HUVECs with WNT2B or GSK3 inhibitor CHIR99021 activates the expression of VEGF. Data represents mean ± SEM. *P* values were calculated by student’s *t*-test or Welch’s *t*-test. Data represents mean ± SEM. *P* values were calculated by ANOVA, Tukey’s test. Data represents mean ± SD. Scale bars, 100 μm. \**P* < .05, \*\**P* < .01, \*\*\**P* < .001, \*\*\*\**P* < .0001. *P* values were calculated by ANOVA, Tukey’s test and student’s *t*-test or Welch’s *t*-test. NC: negative control.

### WNT2B promotes angiogenesis partly through activating VEGF

To identify the WNT ligand responsible for the observed pro-angiogenic effects, we performed qPCR assays to quantify expression of all 19 human WNT ligands in placental villi tissue of each of the three pregnancy types (Figure 4E). Some WNT ligands that have been proposed to activate canonical WNT signaling, such as *WNT2*, *WNT2B* (*WNT13*), *WNT7A*, and *WNT7B*, were significantly up-regulated in IP and REP samples compared with their expression in AEP villi (Figure 4E). Based on the highly significant transcriptional induction in qPCR assays, we selected WNT2 and WNT2B for subsequent analyses of their protein levels in conditioned media, (*P* = 0.0049/0.0030, ANOVA, Tukey’s method). We found that WNT2B was present in high concentrations in IP-conditioned medium compared to WNT2 (100-200 ng/ml vs <10 ng/ml) (Figure 4F). Furthermore, ELISA assays showed that WNT2B was secreted by IP and REP placental villi, but not by AEP villi (Figure 4G), which aligned with the respective differences in the pro-angiogenic potential of their corresponding conditioned media (Figure 3C). These results suggested that WNT2B was likely the ligand responsible for the pro-angiogenic effects of IP-conditioned medium and its upregulation in REP villi tissues implied that this factor could potentially drive vascular development of ectopic placentae associated with prolonged fetal growth in fallopian tubes.

Since WNT2B has not yet been reported as an angiogenic factor in human, we sought to determine whether WNT2B could indeed promote angiogenesis in human cells. To this end, we transfected HEK293FT cells with a plasmid transiently expressing a modified, active human WNT2B (MacDonald et al., 2014) and harvested the conditioned medium for stimulating HUVECs in a sprouting assay. The results indicated that WNT2B-conditioned medium significantly promoted sprout formation, which could be abolished by WIF-1 or sFRP-1 (Figure 4G). To further investigate the mechanism of WNT2B-mediated angiogenesis, we analyzed targets of canonical WNT signaling in HUVECs after treatment with WNT2B-conditioned medium and found that both WNT2B and WNT agonist CHIR99021 significantly induced VEGF expression (Figure 4H). These findings thus established WNT2B as an pro-angiogenic factor for HUVECs and could be responsible for the angiogenic effects of villous conditioned media.

### WNT signaling is required for blood vessel development

The sprouting assay starting from HUVEC cells only measures the angiogenesis effect. To examine whether WNT pathway is also involved in vasculogenesis, we first test whether inhibition of WNT signaling could also influence EC differentiation from human embryonic stem cells that conditionally expressed *ETV2*, which gradually induces EC differentiation (Cakir et al., 2019). To our surprise, only ICG-001 (PRI-724) exhibited strong inhibitory effect on EC differentiation and survival within a 2-day window (Figure 5A). ICG-001 also inhibited sprouting induced by VEGF as demonstrated by HUVEC sprouting (Figure 5B). In contrast, other WNT inhibitors, including XAV-939, and two PORCN inhibitors LGK974 and ETC-159 (Liu et al., 2013; Madan et al., 2016), exhibited strong inhibitory effects on sprouting formation rather than EC differentiation (Figure 5A-B and S5B). Those results, together, suggest that WNT signaling is more likely to be required for angiogenesis rather than vasculogenesis.

**Figure 5.**
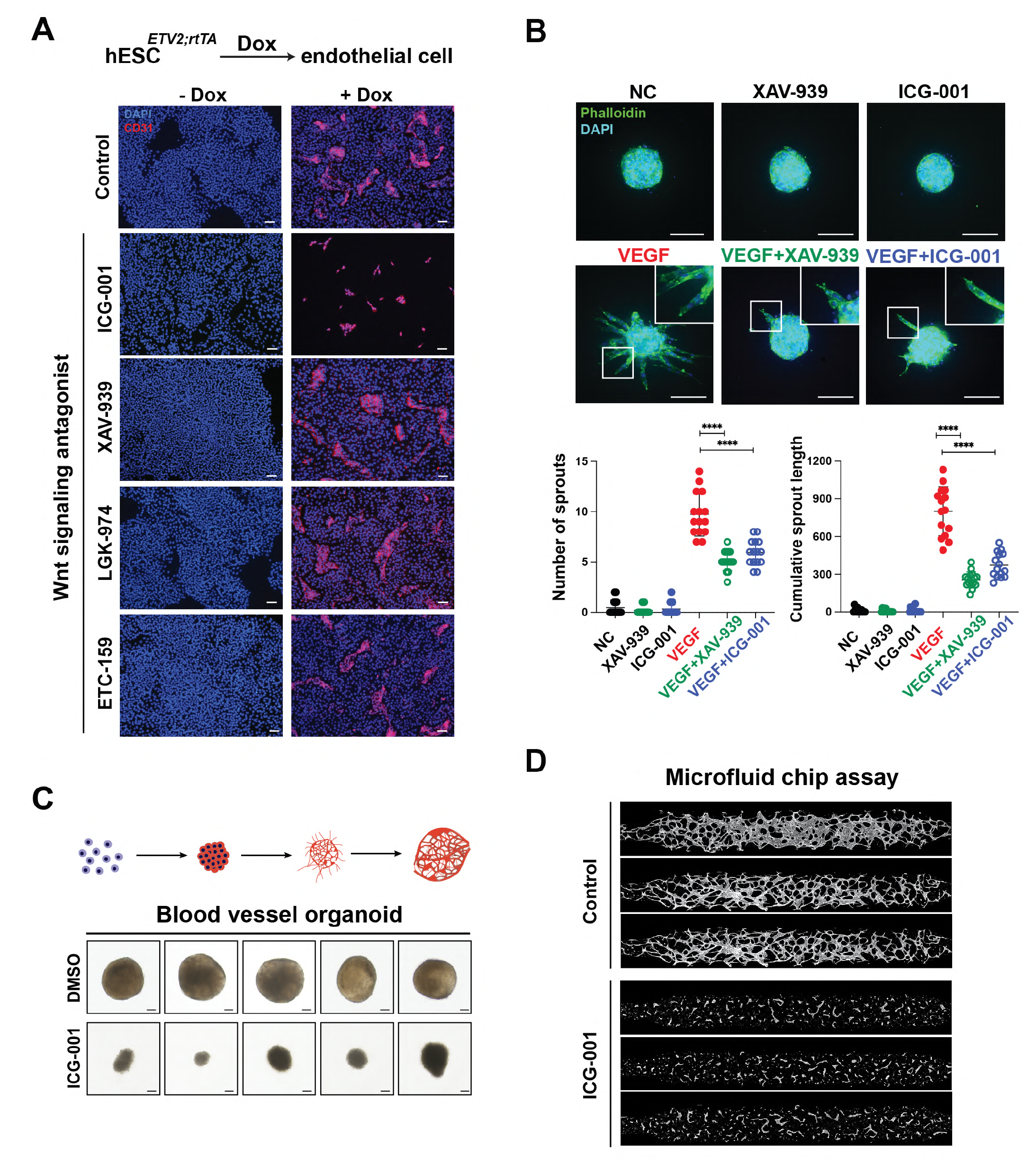
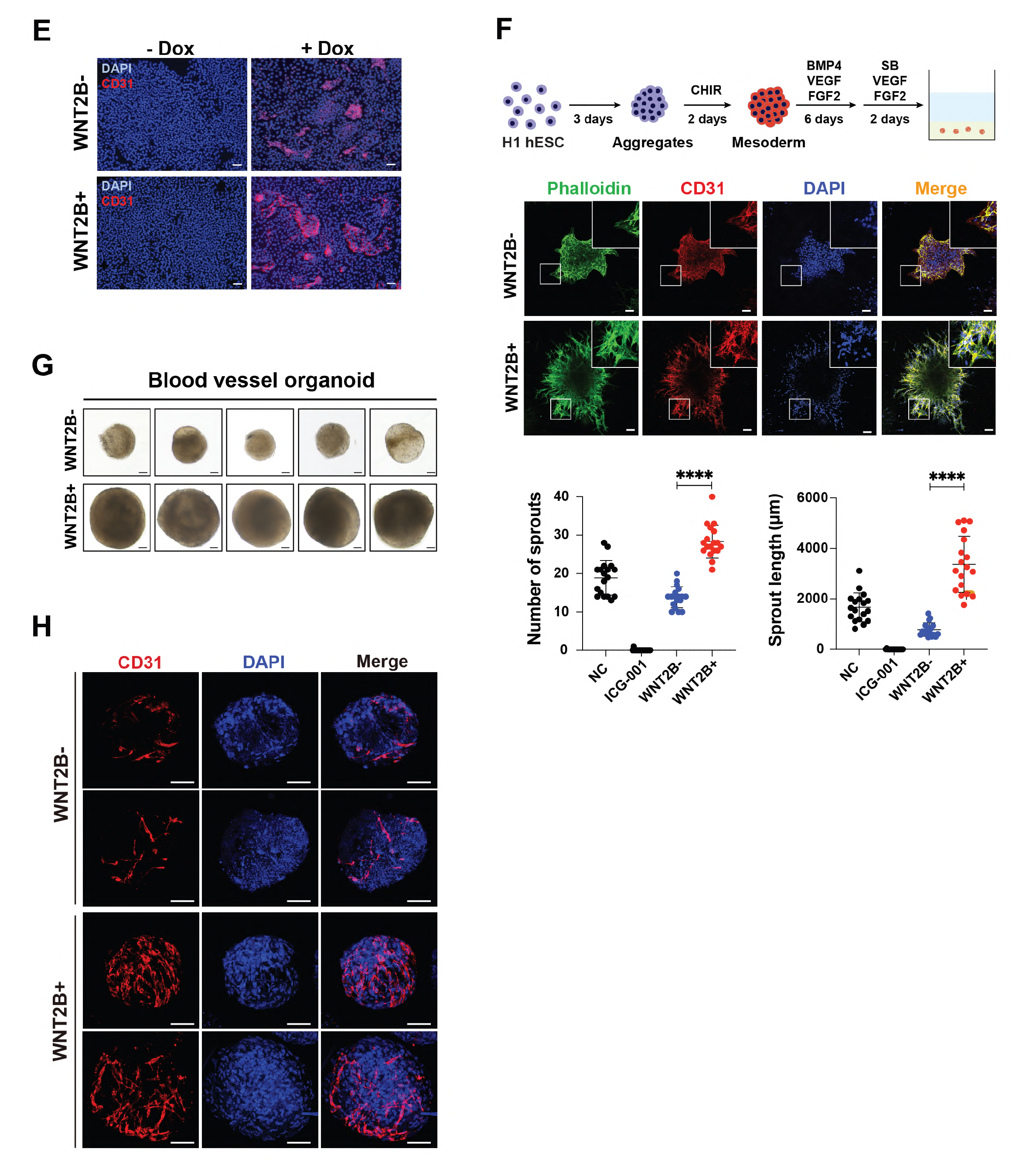
WNT2B promotes blood vessel development on multiple layers. (A) Human embryonic stem cells (hESCs) with ETV2 and rtTA knockin were treated with 2 μg/ml doxycycline for 48 hrs, followed by CD31 staining to examine the endothelial cell fate transition. ICG-001 treatment exhibits pronounced inhibitory effect on endothelial cell transdifferentiation and survival. Scale bars, 100 μm. (B) WNT antagonists ICG-001 (10 μM) and XAV-939 (4 μM) block VEGF-induced sprout formation of HUVECs. (C) Formation of blood vessel organoids were significantly blocked by ICG-001 treatment as evident by decreased sizes of organoids. (D) ICG-001 significantly blocked VEGF-induced vascular network formation of HUVECs in microfluid chip assays. (E) WNT2B promotes the differentiation of hESCs towards endothelial cell fate. 2 μg/ml doxycycline (Dox) were added to the culture medium to induce ETV2 expression in hESCs with 1:1 mixture of conditioned medium in the absence (WNT2B^-^) or presence (WNT2B^+^) of WNT2B for three days. Endothelial differentiation was confirmed by CD31 staining. (F-H) WNT2B promotes vascular specification and vascular network development in blood vessel organoids. WNT2B-conditioned and control media were supplemented with differentiation factors and added at the time cell aggregates were embedded in 3D gel matrices and differentiated for 5 days for sprout measurement (F). Cell aggregates primed for EC specification were then transferred to suspension culture for further differentiation to examine the vascular network formation by CD31 immunostaining (G, H). Scale bars, 100 μm. Data represents mean ± SD. \**P* < .05, \*\**P* < .01, \*\*\**P* < .001, \*\*\*\**P* < .0001. *P* values were calculated by ANOVA, Tukey’s test. NC, negative control. CM, conditioned medium.

Recently developed blood vessel organoids derived from human pluripotent stem cells recapitulate the continuous process of mesoderm development, vascular specification and network expansion (Wimmer et al., 2019). We therefore tested whether WNT signaling is also required for the development of blood vessel organoids. Treatment of ICG-001 greatly shrank the size of blood vessel organoids (Figure 5C). We further validated the role of WNT2B in vascular network formation and expansion in an *in vitro* microfluidic chip assay which provides unprecedented advantage for evaluating complex vascular network than other *in vitro* assays (Kim et al., 2013). The results showed that inhibition of WNT signaling by ICG-001 dramatically disrupted the vascular network formation in the microfluidic chips (Figure 5D). These results suggested that inhibition of β-catenin/CBP compromises the vasculogenesis, angiogenesis and vascular network formation.

### WNT2B promotes blood vessel formation at vasculogenesis, angiogenesis, and network expansion levels

Since WNT2B promotes sprouting of HUVEC (Figure 4F), we next examined whether WNT2B also promotes EC differentiation. Adding WNT2B^+^ conditioned medium to the differentiation medium significantly increased CD31^+^ cell numbers in this transdifferentiation system as compared with WNT2B^-^ conditioned medium (Figure 5E). To further study at which stages WNT2B influences human blood vessel development, we dissected the formation of blood vessel organoids into vascular specification (cultured in 3D hydrogel) and vascular network formation (cultured in suspension) stages (Wimmer et al., 2019). Treatment of WNT2B^+^ conditioned medium greatly increased the number and length of CD31^+^ sprouts of blood vessel organoids at the vascular specification stage (Figure 5F). Importantly, we found WNT2B-conditioned medium not only significantly increased the size of blood vessel organoids, but also largely promoted the formation of vascular network in blood vessel organoids (Figure 5G-H). Based on these results, we conclude that WNT2B likely promotes vascularization at vasculogenesis, angiogenesis, and vascular network expansion levels.

### WNT2B secreted by syncytiotrophoblasts is greatly decreased in AEP condition

We next try to find the source of WNT2B in placental villi. Within the range of gestational ages analyzed here (40-64 days), the placental villi developed into the tertiary stage, forming dendritic chorionic villi in which each branch consisted of blood vessels, central stromal cells, the outer-most syncytiotrophoblast (STB) layer, a lower villous cytotrophoblast (vCTB) layer, and extravillous trophoblasts (EVTs) located at tips of anchoring villi (Figure 1B) (Baergen, 2005). Since the placentae develop to different extents at similar gestational ages in REP and AEP, we further examined whether there were morphological or cellular differences among the chorionic villi of IP, AEP, and REP patients. Surprisingly, in all three types of placental villi, morphologically and structurally similar trophoblast shells (the STB and vCTB layers) surrounded the trophoblastic columns, as shown by histological and CK7 staining (Figure 6A-B). Furthermore, we observed similar extracellular and intracellular patterns of syndecan 1 (SDC-1), an extracellular marker of STBs, across all three types of placental villi, suggesting comparable STB maturation (Figure 6B). Immunostaining of TEA Domain Transcription Factor 4 (TEAD4), a vCTB marker, also revealed that the number and arrangement of vCTBs was similar for all three types of placental villi (Figure 6B). Notably, we found that AEP and REP villi had larger diameter of trophoblastic columns compared with those of IP patients (Figure 6C). However, similar relative ratios of proliferative (KI67^+^TEAD4^+^) to quiescent (KI67^-^TEAD4^+^) vCTBs implied that proliferation of stem cell populations was also similar in the three types of placental villi (Figure 6C) (Liu et al., 2018). Excluding differences in trophoblastic column diameter, histological and immunological analyses showed only minor, if any, differences in morphology, arrangement, trophoblast differentiation, and stem cell activity in the trophoblasts of placental villi from the three types of pregnancies.

**Figure 6.**
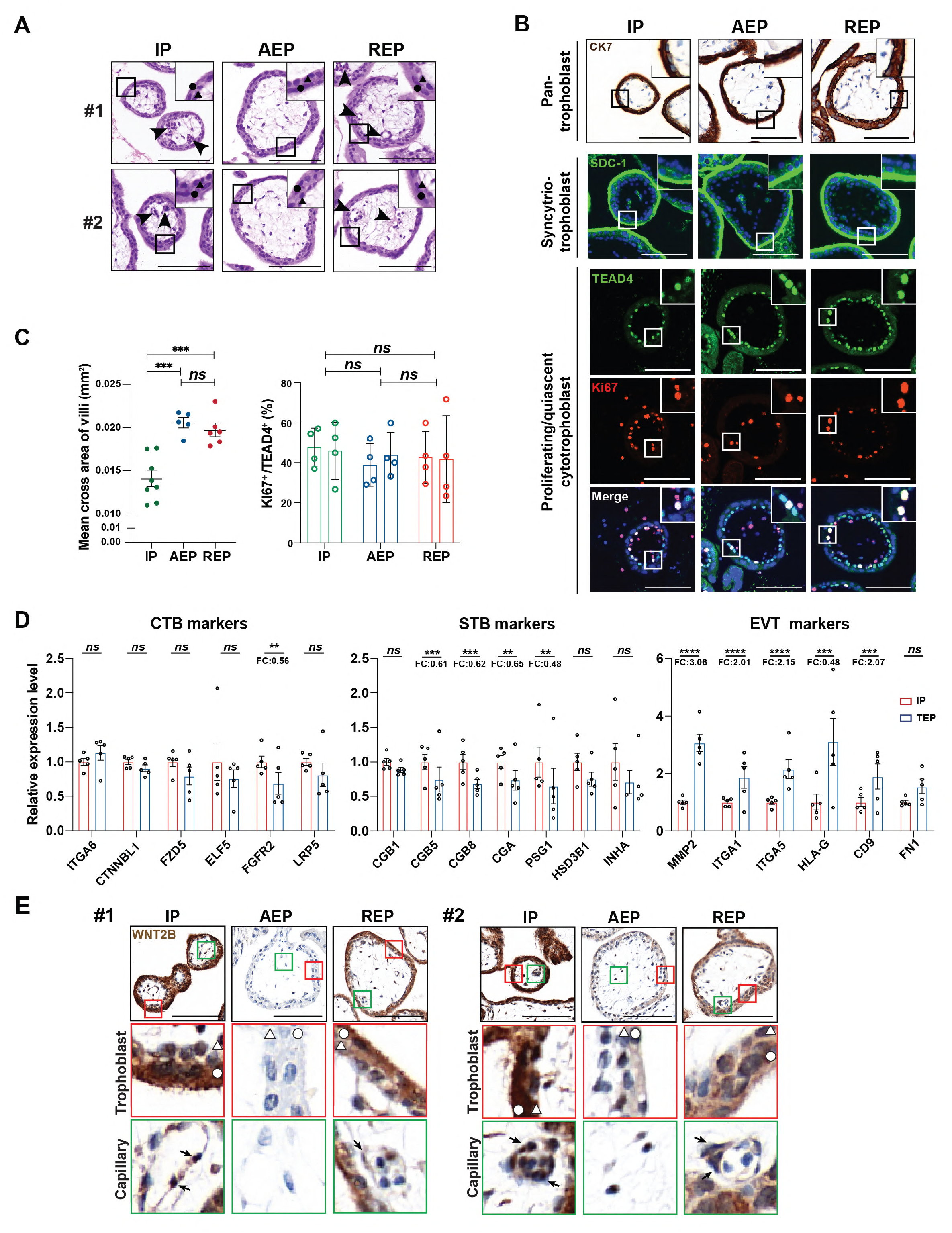
WNT2B are secreted from syncytiotrophoblasts in placental villi. (A-C) Comparison of trophoblasts in IP, AEP, and REP pregnancies. (A) H&E staining of placental villi cross sections from three types of pregnancies. Two clinical samples are shown for each type of pregnancy. Arrows, blood vessels. Scale bars, 100 μm. (B) Representative images of immunostaining of the trophoblast marker CK7, STB marker SDC1, vCTB marker TEAD4, and proliferative marker KI67 in placental villi of three pregnancies. Similar extracellular and intracellular patterns of SDC1 and similar number and arrangement of proliferative and non-proliferative vCTBs were observed in three types of pregnancy. Scale bars, 100 μm. (C) Left, quantification of the mean cross area of proliferative vCTBs of villi from IP (n=8), AEP (n=5) and REP (n=6). For each sample, data was collected from at least eight independent fields of views. The graph shows the mean ± SEM. Right, quantification of the ratio of proliferative vCTBs of villi from IP (n=2), AEP (n=2) and REP (n=2). Similar proliferation activities of the stem cell population were observed in three types of placental villi. (D) Expression of trophoblast-specific genes in IP (n=5) and TEP (n=5) placental villi suggests minor differences in CTB and STB marker expression, but significant upregulation of EVT markers in the TEP condition. Log-transformed, normalized expression levels for selected typic markers of three trophoblast subsets are presented as mean ± SEM. \*\**P* < .01, \*\*\**P* < .001, \*\*\*\**P* < .0001. *P* values were calculated by student’s *t*-test or Welch’s *t*-test. (E) WNT2B is produced by trophoblasts of IP and REP placental villi, not AEP ones, as revealed by immunohistochemical staining for WNT2B. Two clinical samples (#1 and #2) for each type of pregnancy were presented. Red rectangle, trophoblasts. Green rectangle, blood vessels. Triangles, vCTB. Circles, STB. Scale bars, 100 μm.

We next compared the expression of trophoblast markers between IP and TEP villi and found only minor differences in some markers of vCTB and STB (Figure 6D). These findings indicated that implantation in the fallopian tubes did not substantially compromise the self-renewal of vCTBs and their differentiation potential toward STBs. In contrast, we did observe significant upregulation of EVT markers (*e.g., MMP2, ITGA1, ITGA5* and *HLA-G*) in placental villi of TEP (Figure 6D), which could be due to the relatively hypoxic condition in the fallopian tube (Chakraborty et al., 2011). The skewed differentiation of vCTB towards invasive trophoblasts (ETVs) was also consistent with dysregulation of the ECM, cell adhesion molecules, and secretory metallopeptidases (Figure 2G and S3E).

We next performed immunohistochemical staining of IP, REP, and AEP placental villi with anti-WNT2B antibody to identify the cell population responsible for WNT2B production. The results indicated that WNT2B was strongly expressed by trophoblasts (primarily STBs, with slightly lower expression in vCTBs and mild expression in ECs) in the trophoblastic columns of IP and REP villi (Figure 6E). Consistently, the expression level of WNT2B was largely decreased in AEP (Figure 6E). Those results suggested that the aberrant elevation of WNT2B level in the trophoblasts embedded in fallopian tubes could lead to excessive vascularization and eventually the rupture of fallopian tubes.

### Organoid cocultures reveal the essential role of trophoblast-secreted WNT2B in promoting vascularization

Since WNT2B is secreted from trophoblasts, we aim to develop a composite cell culture model to study how trophoblasts interact with endothelial cells/ endothelial progenitor cells to influence vascular specification (Figure 7A). We first derived two trophoblast stem cell (TSC) lines from human blastocysts (Figure 7B) (Okae et al., 2018). After establishing control and WNT2B-silenced TSC lines (Figure 7C-D), digested TSCs and blood vessel organoids at the stage of vascular lineage induction were simultaneously embedded in corresponding hydrogels within the Boyden and the bottom chamber, respectively, and cultured with TOM (trophoblast organoids) and blood vessel organoid medium (Turco et al., 2018; Wimmer et al., 2019). After six days, the matured trophoblast organoids were transferred and cultured with new blood vessel organoids (also at the vascular lineage induction stage) to investigate the interaction between these two organoids (Figure 7A). Interestingly, coculturing with trophoblast organoids did promote the sprouting of blood vessel organoids as compared with the control (hydrogel alone) (Figure 7E), consistent with the pro-angiogenic effect of conditioned media from placental villi (Figure 3C). Importantly, we were able to demonstrate that the pro-angiogenic effect of trophoblast organoids are dependent on WNT2B in this organoid coculture model, since knockdown of WNT2B significantly compromised the sprouting of blood vessel organoids (Figure 7E). Together with results from other assays, this composite culturing of trophoblast organoids and blood vessel organoids recapitulates the interaction between trophoblasts and blood vessel lineages and confirms the causal relationship between WNT2B and trophoblast-promoted vascularization.

**Figure 7.**
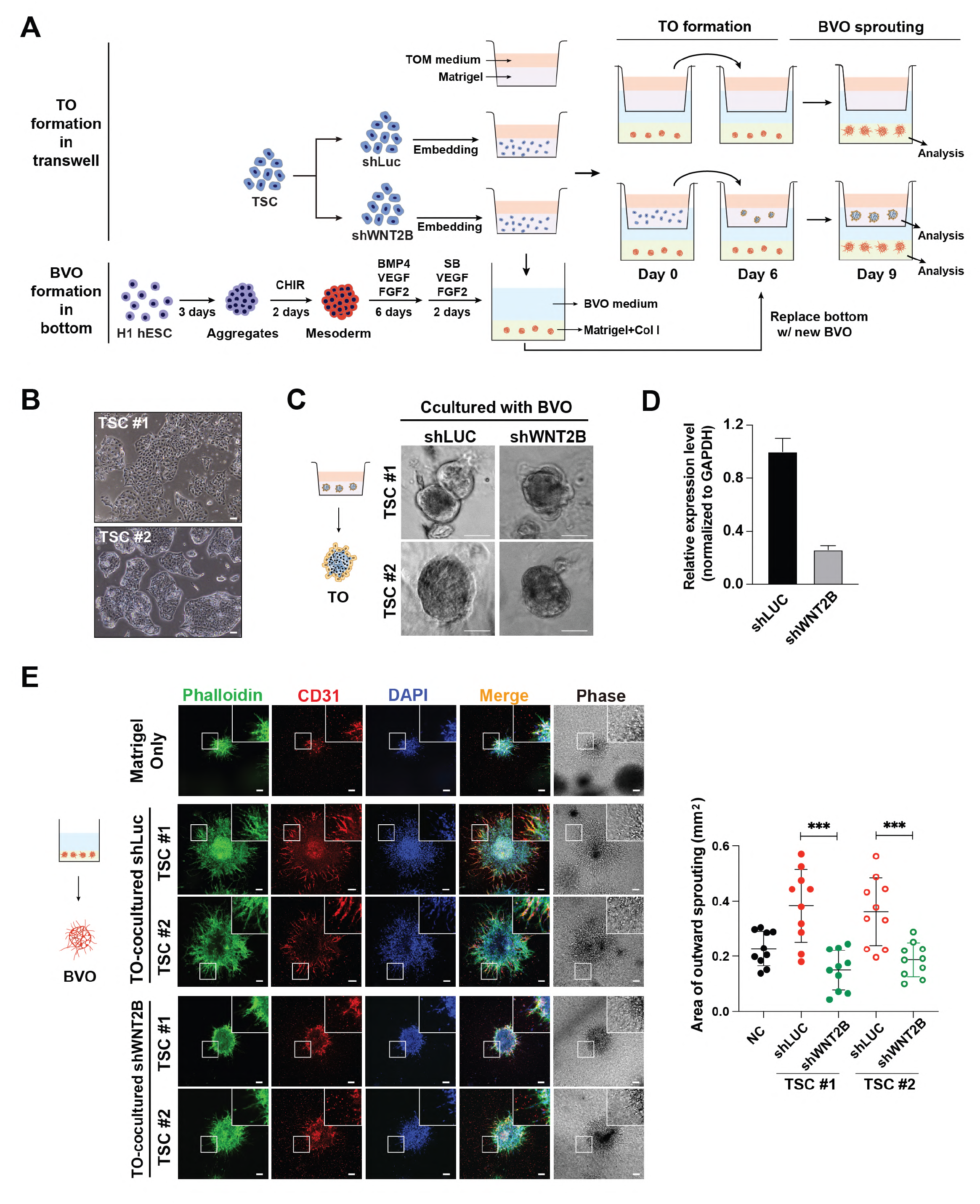
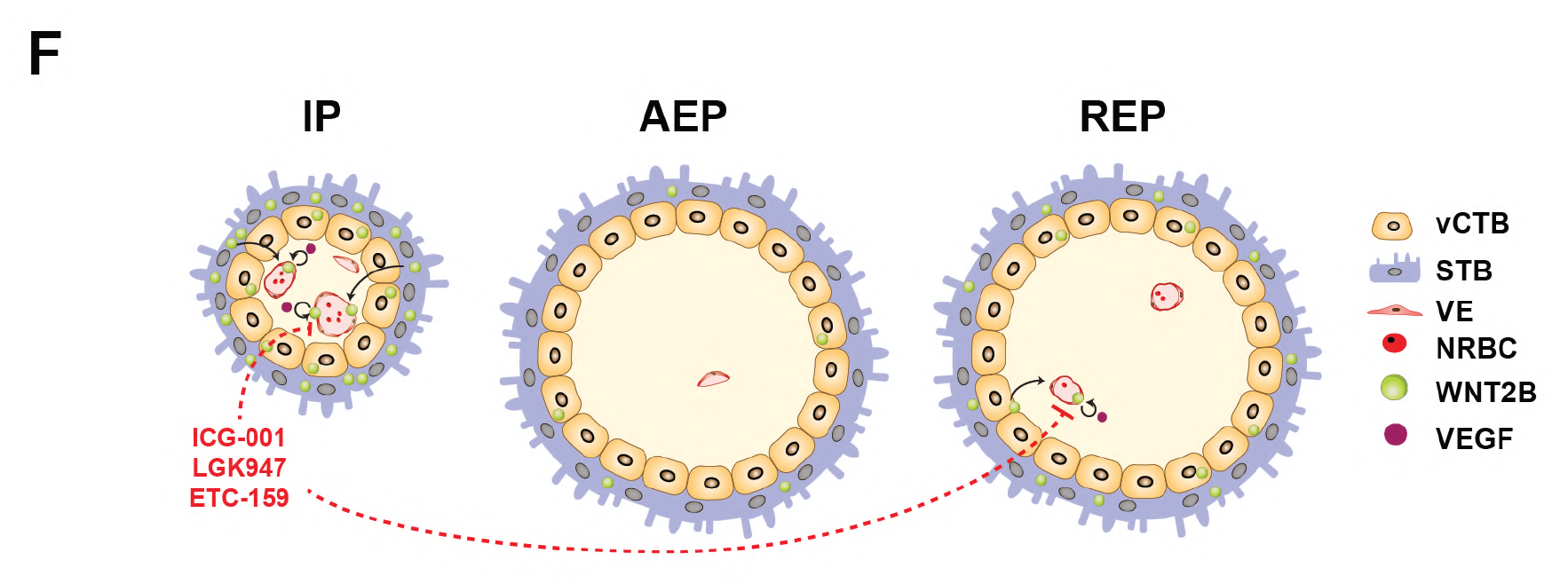
The organoid coculture model revealed the role of WNT2B in human vascularization. (A) Schematic illustration of the coculture system containing trophoblast organoids (TOs) and blood vessel organoids (BVOs). Trophoblast organoids were formed in the presence of cell aggregates containing specified endothelial cells/projectors. Matured trophoblast organoids were then transferred and cocultured with another set of cell aggregates to investigate the influence of TOs on BVO development. (B) Two independent trophoblast stem cell lines derived from human blastocysts exhibited normal trophoblast stem cell morphologies. Scale bars, 100 μm. (C) Morphologies of TOs formed in the coculture system, in which WNT2B was significantly knocked down by shRNAs (D). Scale bars, 50 μm. (E) TOs significantly promoted the development of BVOs in a WNT2B-dependent manner. CD31 immunostaining (left) and statistics (right) of the sprouts of BVO after coculturing with Matrigel alone or TO for 3 days. Scale bars, 100 μm. (F) The schematic illustration and summary of three pregnancy types of villi and intravillous capillaries. Data represents mean ± SD. \**P* < .05, \*\**P* < .01, \*\*\**P* < .001, \*\*\*\**P* < .0001. *P* values were calculated by ANOVA, Tukey’s test.

### The microenvironment of fallopian tubes modulates WNT2B expression

In light of these results, we proposed intrinsic and extrinsic hypotheses to explain why WNT2B is differentially produced in REP but not AEP placental villi. In the intrinsic hypothesis, AEP and REP trophoblasts harbor distinct genetic or epigenetic mutations that result in aberrant WNT2B expression. Alternatively, in the extrinsic hypothesis, the AEP and REP tubal microenvironments exhibit different crosstalk signals with trophoblasts, resulting in an indirect influence on WNT2B secretion. To test these two possibilities, we derived trophoblast organoids (TOs) from IP and TEP trophoblasts using previously published methods (Figure S6A) (Turco et al., 2018). To our surprise, IP- and TEP-derived TOs, generated from isolated villous stem cells (vCTB), and cultured under the same *in vitro* conditions, exhibited similar morphologies, transcriptome patterns, as well as WNT2B and other trophoblast marker expression levels (Figure S6B-D). Interestingly, hCG expression levels, the diagnostic marker which distinguishes AEP and REP, were indistinguishable between IP and TEP TOs (Figure S6E). Based on these results, it appears likely that extrinsic factors in the microenvironment affect WNT2B expression via communication with trophoblasts.

In summary, by employing multiple assays, we identified a novel WNT-dependent mechanism for intravillous angiogenesis during pregnancy (Figure 7F). This WNT-promoted angiogenesis is attributable to WNT2B secreted by the trophoblast shells of placental villi. WNT2B-regulated development of the villous vasculature is crucial for establishing fetal circulation. In TEP, ectopic implantation in fallopian tubes frequently leads to AEP, in which low WNT2B expression restricts angiogenesis and therefore further fetal growth. Alternatively, aberrant expression of WNT2B in certain conditions of TEP could result in extensive vascular formation in placental villi, leading to excessive fetal growth and tubal rupture.

## Discussion

Ectopic pregnancy (EP) is a critical pregnancy complication and a major cause of pregnancy-related death. However, the pathological factors that lead to the rupture phenotype remain unknown and controversial. With particular care to exclude confounding factors, in this study we interrogated clinical villi samples from intrauterine as well as ruptured and abortive tubal ectopic pregnancies, which revealed the development of excessive intravillous vascularization in ruptured EP. A recent, large-scale knockout study in mice revealed the strong correlation between placental defects and embryonic developmental abnormalities (Perez-Garcia et al., 2018). Notably, a number of knockouts showed detrimental impacts on the growth and organization of fetal and maternal blood conduits within the mouse labyrinth layer, the functional analog of human placental villi (Perez-Garcia et al., 2018). These findings, together with ours, suggest a central role for intravillous angiogenesis in both normal embryonic development as well as in pregnancy complications in humans.

Vascular remodeling during pregnancy can be viewed from either a maternal or fetal perspective. From the maternal perspective, the angiogenesis that occurs during decidualization and the remodeling of spiral arteries together ensure a sufficient maternal blood supply. In contrast to maternal angiogenesis, less is known about fetal intravillous angiogenesis, which in fact represents an essential factor in the development of adequate fetal circulation. Previous studies have identified PROK1 as a factor that promotes intravillous angiogenesis potentially associated with pregnancy complications (Alfaidy et al., 2014). Interestingly, we identified trophoblast-secreted WNT2B as both a vasculogenic and an angiogenic factor required for initiating fetal intravillous vascularization in both IP and REP. WNT2 and WNT2B, which share 67% amino acid homology, are both described as canonical WNT ligands. However, in mice, the *wnt2b* knockout phenotype is largely indiscernible from that of wild type, except for distinct olfactory bulb traits (Goss et al., 2009; Tsukiyama and Yamaguchi, 2012). In contrast, *wnt2* knockout resulted in strong vasculature defects in placentae and multiple organs, suggesting a major contribution to placental development that more closely resembles our findings of WNT2B function in humans (Goss et al., 2009; Monkley et al., 1996). The fact that WNT2 levels did not significantly differ between IP- and EP-conditioned media suggests that WNT2B rather than WNT2 plays the major role in promoting intravillous angiogenesis in humans (Schmidt et al., 2015). Interestingly, both WNT2 and WNT2B have been reported in other human pregnancy complications (Li et al., 2017; Zhang et al., 2018). It’s worthy to investigate the role of WNT-dependent angiogenesis in other pathological contexts.

The WNT signaling pathway has been shown in humans/rodents to promote or maintain angiogenesis in multiple organs as well as in tumors (Wang et al., 2019). Our data showed that Wnt stimulate VEGF expression in EC, which is consistent with previous results that VEGF as a downstream target of the WNT signal cascade (Zhang et al., 2001; Zhang et al., 2016). Previous results showed that WNT signaling is required for VEGF-induced endothelial differentiation from mesenchymal stem cells (Zhang et al., 2016). Our study further demonstrates that WNT2B can stimulate EC differentiation from pluripotent stem cells in the absence of VEGF. In addition, by employing the blood vessel organoid model, we showed the WNT2B could also enhances vascular network formation. Future investigations will help determining whether those three vascular formation processes promoted by WNT2B share the same molecular mechanism.

In summary, our findings employed organoid-based models to reveal the molecular basis underlying the etiology of REP and shed important insight on the potential of developing a novel therapeutic strategies. First, WNT2B, the WNT signal pathway, or vasculogenic structures in placental villi could be adopted as diagnostic markers in monitoring for REP or in the selection of appropriate therapeutic strategies, to exclude AEP for expectant or medical therapy. Second, antagonizing WNT2B, WNT signal pathway, or angiogenic factors with neutralizing antibodies or inhibitors could be a viable approach to inhibit or diminish ectopic fetal growth in fallopian tubes, thereby alleviating symptoms. Third, since WNT2B overproduction likely results from cues in the tubal wall microenvironment, antagonizing the factors or cell-cell interactions that induce WNT2B expression could also represent a promising avenue for therapeutic development. Clinical explants and organoid-based models employed in this study will thus facilitate exploring the mechanisms of WNT2B production and WNT-dependent vascular development to improve our understanding of the intricate crosstalk at the maternal-fetal interface in eutopic and ectopic pregnancies.

## Acknowledgments

This research was supported by National Key R&D Program of China (2020YFA0710800, 2021YFC2700100, and 2018YFC1004900), Key R&D Program of Zhejiang (2021C03G2013079), Shanghai Science and Technology Commission (19JC1414200), National Natural Science Foundation of China (J.Z., 82171667; C.L., 31871487; D.Z., 81801400 and W.X., 81901503), Strategic Collaborative Research Program of the Ferring Institute of Reproductive Medicine (FIRMA200506), and the ShanghaiTech University start-up fund. We thank the Multi-Omics Core Facility (MOCF), Molecular Imaging Core Facility (MICF), and Molecular and Cell Biology Core Facility (MCBCF) at the School of Life Science and Technology, ShanghaiTech University for providing technical support. We would like to thank statistician of Li Xie for expert advice on statistical analyses. We apologize for critical works that are not cited due to space constraints.

## Author Contributions

Contribution: X.Z., Z.Z. designed and performed research and analyzed data; S.S. and Y.L. performed research; Q.Z., D.Z. and W.X. collected clinical samples; Q.Y. analyzed the sequencing data; L.J and Y.X generated the ETV2-knockin human embryonic stem cells; C.L and J.Z. designed and performed research, analyzed data, and wrote the manuscript.

## Methods

### 3D reconstruction of placental villous vascular networks

The protocol of clarification and 3D reconstruction of immunolabeled placental villous vascular network was from the Merz G et al (Merz et al., 2018). Villous trees were incubated with anti-CK7 and anti-CD31 antibodies. Samples were treated with HISTO-1 (Visikol) and HISTO-2 (Visikol) for 4h at RT for clarification. The clarified tissues were placed in HISTO-2, put in the Sykes-Moore chamber (BellCo Glass) and scanned using Leica SP8 confocal microscope.

### Explant of tissues from maternal-fetal interfaces and ELISA

Explants were cultured according to a previously published protocol with minor modifications (James et al., 2006). Briefly, maternal-fetal interfaces were surgically dissected. Villous tips or maternal tissues (uterine endometria or fallopian tube walls) were separated, weighted, and placed in the center of wells of 48-well plates. The villous explants were incubated in basal media consisting of advanced DMEM/F12 (Life Technologies, #12634-010) containing 1% Glutamax (Life Technologies, #35050-061), 100 μg/ml streptomycin and 100 U/ml penicillin (Life Technologies, #15140-122). Human endometrial and fallopian tube explants were also cultured in basal media. The explants were strictly cultured in 5 μl medium volume per mg wet weight. The conditional medium was collected every 24 hours for 2 days.

Conditioned media were collected from explant cultures, centrifuged at 3000 rpm to remove debris and stored at −80°C until use. The target protein levels in conditional media were quantified using ELISA following manufacturer’s instructions. Vascular endothelial growth factor (VEGF) ELISA kits were purchased from Absin, Shanghai (#abs510008-96T); WNT2 and WNT2B were measured using ELISA kit from FineTest, Wuhai (#EH2251 and #EH2250). Each sample with three replications.

### RNA-seq analyses

Total RNA was extracted from the chorionic villi samples or placental villus organoids using Trizol (Invitrogen, #15596026) according to manufacturer’s instruction. The RNA purity was checked using the kaiaoK5500 spectrophotometer (Kaiao, Beijing, China). The RNA integrity and concentration were assessed on Bioanalyzer 2100 system (Agilent Technologies, CA, USA) using RNA Nano 6000 Assay Kit. All primary samples have RNA integrity number equivalent (RINe) values > 8.5. Sequencing libraries were generated using NEBNext Ultra RNA Library Prep Kit for Illumina (#E7530L, NEB, USA) following the manufacturer’s recommendations and index codes were added to attribute sequences to each sample. Briefly, mRNA was purified using poly-T oligo-attached magnetic beads, and then fragmented and reverse transcribed into double-stranded complementary DNA. The library fragments were purified with QiaQuick PCR kits and elution with EB buffer, then terminal repair, A-tailing and adapter added were implemented. The aimed products were retrieved and PCR was performed, then the library was completed. RNA concentration of library was measured using Qubit RNA Assay Kit in Qubit 3.0 to preliminary quantify and then dilute to 1ng/μl. Insert size was assessed using the Agilent Bioanalyzer 2100 system (Agilent Technologies, CA, USA), and accurated quantification using StepOnePlus Real-Time PCR System (Library valid concentration > 10 nM). The libraries were then sequenced on the Illumina HiSeq PE Cluster Kit v4-cBot-HS (Illumina) with 150-bp paired-end reads (GSE183319 and GSE185119). The reference genomes (Homo sapiens GRCh38) and the annotation file were downloaded from ENSEMBL database (http://www.ensembl.org/index.html).

Bowtie2 v2.2.3 was used for building the genome index, and Clean Data was then aligned to the reference genome using HISAT2 v2.1.0 (Kim et al., 2015). For female samples, the Y chromosome was excluded from the reference genome. The read count for each gene in each sample was counted by HTSeq v0.6.0, and then calculated to estimate the expression level of genes by DEseq2 (Anders and Huber, 2010; Love et al., 2014). Differentially expressed genes were identified using DESeq2 (Anders and Huber, 2010; Love et al., 2014).

A fold change ≥1.5 and a P value < 0.05 were considered to indicate the differentially expressed genes (DEGs) (Table S4). The obtained DEGs were visualized in a volcano and heatmap plots with gplots (Warnes et al., 2005) and ggplot2 (Wickham, 2016) R Bioconductor packages [http://www.r-project.org/]. Furthermore, more in-depth analyses based on the DEGs, including GO enrichment analysis, KEGG pathway enrichment analysis, and Gene Set Enrichment Analysis (GSEA).

### Tubal network formation angiogenesis assay

Examining the formation of tube-like structures. Once HUVECs reaches 70-80% confluence, the cells were serum starved for 4 hours so that it performs the tube formation assay better (Ko and Lung, 2012). Briefly, distributing 50 μl growth factor-reduced Matrigel (Corning, #354230) per well of the pre-chilled 96-well plate on ice and incubating at 37 °C for 30 min to polymerization. 1*105 HUVECs were resuspended in the conditional medium (100 μl) and seeded onto the 96-well plates precoated with Matrigel (Corning, #354230). After 24 hours, tube formation was quantified by measuring the cumulative tube length, loop count, loop area, and branch count with FIJI (Tetzlaff and Fischer, 2018).

### Sprouting angiogenesis assay

To examine capillary sprouting, HUVECs were used to generate spheroids at the density of 800 cells/spheroid in-gel as previously described (Zahra et al., 2019). The spheroids were cultured in basal media or conditioned media from explants or WNT2B-overexpressing HEK293T cells containing one or multiple following factors. At the first day for spheroids formation, HUVECs were suspended in the EGM2/methylcellulose medium (40 cells/μl) and distributed in 20 μl hanging drops on the lid of 10 cm dishes and incubated at 37°C for 24 h. At the second day, spheroids were harvested and embedded in the methylcellulose/collagen type I gel mix, followed by incubation for 30 min at 37°C for polymerization. Subsequently, spheroids were cultured in basal medium only or with factors at 37°C for 24 h. Statistical analyses of HUVEC sprouts were performed by randomly selecting 15 spheroids, measuring the number of sprouts per spheroid (Tetzlaff and Fischer, 2018). Each experiment was repeated at least two times.

### WNT2B conditioned medium

The WNT2B conditioned medium was generated via transfecting HEK293FT cells with the pcDNA3.1-WNT2B plasmid (synthesized by Tsingke Biological Technology, China) (MacDonald et al., 2014). Control HECK293FT cells were transfected with the empty pcDNA3.1 vector. After cells grew to confluence in DMEM/F12 (10% FBS), media was replaced with basal medium for 24 hours before collection.

### Endothelial cell differentiation

Human H9 embryonic stem cells with *tetO-ETV2* and *rtTA* knock-in (the detailed editing strategy will be described in other studies) were used to generate endothelial cells. Briefly, cultured stem cell colonies were directly treated with 2 μg/ml doxycycline in mTeSR-1 medium for 48 hrs. The appearance of endothelial cells was then examined by CD31 immunostaining.

### Microfluidic chip assay

Microfluidic chips were designed and fabricated according to the previous study (Kim et al., 2013). In brief, HUVEC and HPF were trypsinized and resuspended in 5 mg/mL fibrinogen (Sigma, F8630), supplemented with 0.30 U/mL aprotinin (Sigma, #A1153) at a density of 12E6 cell/mL (HUVEC) and 4E6 cell/mL (HPF). The 2X mixture was then mixed with 2 U/mL thrombin (Sigma, #T4648) at a 1:1 (v:v) ratio. The central channel and side channel were filled with HUVEC cell solutions and HPF cell solutions respectively and left to clot at room temperature for 5 minutes. Chips were then placed in a humid chamber and given basic endothelial growth medium 2 with and treated with 10 µM ICG-001 for five days and changed medium every two days during this time. At day 5, cells were fixed by introducing 4% PFA into the medium channel for 1 hour at room temperature and then staining with phalloidin.

### Blood vessel organoid formation assay

Blood vessel organoids were generated from human H1 embryonic stem cell following Wimmer et al except EGM-2 was used instead of StemPro-34 medium (Wimmer et al., 2019). For mifepristone treatment, the drug was added at the time inducing vascular lineage differentiation (day 3). For WNT2B treatment, WNT2B-conditioned EGM-2 medium was added at the time cell aggregates were embedded into 3D matrix (day 5). Organoids were fixed at day 8-9 and immunostained with CD31 antibodies.

### Trophoblast organoid culture

Derivation of human trophoblast organoids (TOs) was based on the previously published method (Turco et al., 2018). TOs in matrigel drops were carefully overlaid with 250 μl trophoblast organoid medium (TOM) consisting of Advanced DMEM/F12 (Life Technologies) supplemented with 1×N2, 1×B27, 2 mM Glutamax, 100 μg/ml streptomycin and 100 U/ml penicillin, 1.25 mM N-Acetyl-L-cysteine, 500 nM A 83-01, 1.5 μM CHIR99021, 50 ng/ml recombinant human EGF, 80 ng/ml recombinant human R-spondin 1, 100 ng/ml recombinant human FGF2, 50 ng/mL recombinant human HGF, 10 mM nicotinamide, 2 μM Y-27632, and 2.5 μM prostaglandin E2 (PGE2). The baseline characteristics are provided in the Supplemental Table 3.

### RNA-seq analyses

RNA-seq reads (GSE183319 and GSE185119) were aligned to the *Homo sapiens* GRCh38 reference genome. The read counts for each gene in each sample was counted by HTSeq v0.6.0, and then calculated to estimate the expression level of genes by DEseq2 (Anders and Huber, 2010; Love et al., 2014).

### Statistics

The student’s *t*-test, Welch’s *t* test and one-way ANOVA were used to determine the statistical significance. Tukey’s method was used in one-way ANOVA to make multiple comparisons. Continuous variables are presented as mean ± standard deviation (SD)/standard error of mean (SEM). The linear regressions of measured parameters were calculated with ages and gestational days as independent variables.

### Study Approval

All human tissues used in the study were obtained through the International Peace Maternity and Child Health Hospital (IPMCH) in Shanghai, China and taken with the written informed consent from Chinese pregnant women. Human blastocysts used for trophoblast stem cell derivation were obtained through the Women’s Hospital, Zhejiang University School of Medicine. The study was approved by the institutional ethics committee of the IPMCH (GKLW201909) and Zhejiang Women’s Hospital (SC-2019001). Details about distribution of samples according to baseline characteristics are provided in the Figure S1A and Table S1-3.

