## Supplemental Information for "Trophoblast and blood vessel organoid cultures recapitulate the role of WNT2B in promoting intravillous vascularization in human intrauterine and ectopic pregnancy"

**Cell culture**

HEK293FT cells were cultured in DMEM (Life Technologies, #C11965500BT) containing 10% FBS (Life Technologies, #16000-044), and 100 μg/ml streptomycin and 100 U/ml penicillin (Life Technologies, #15140-122). Human umbilical vein endothelial cells (HUVECs) were isolated from umbilical cords of normal full-term natural delivery as previously described (Crampton et al., 2007). The pregnancy of normal full-term natural delivery (39 weeks + 2 days) is 29 years old without any pregnancy complications. In brief, the vein of the umbilical cord was washed with cold phosphate-buffered saline (PBS), infused and incubated with type I collagenase (1 mg/ml, Life Technologies, #17100-017) at 37°C for 15 min for three times. Resultant isolated cells were grown in endothelial growth medium 2 (EGM-2) (Lonza, #CC-3124). EGM-2 medium was replaced every 2 days and HUVECs were passaged every 3 to 4 days depending on the density. HUVECs at passage 2–5 were used in the present study.

**Case collection**

Intrauterine placental villi for histological and immunological analyses, tissue clearing, explant culture, RT-qPCR, and ELISA were collected from 41, for RNA-seq was collected from 5, and for culturing trophoblastic organoids (TOs) from 5 normal and voluntarily terminated intrauterine pregnancies (IP) that confirmed by ultrasonography combined with blood test or urine pregnancy test. Tissue samples were collected immediately after surgically resected chorionic villi and fallopian tube/uterine endometrium. Then, transported tissues from operation room to laboratory within 15 minutes. For ectopic pregnant placental villi (including AEP, REP that contains 34 and 37, respectively), the tubal implantation was first diagnosed by ultrasonography, followed by confirmation and removal by salpingectomy. All TEP samples were collected without receiving methotrexate treatment. All clinical placental villous samples obtained from Chinese pregnant women at the Department of Obstetrics and Gynecology of the IPMCH from October 2019 to June 2021 and divided into IP (n=41), AEP (n=34), and REP (n=28) groups for histological and immunological analyses, explant culture, and ELISA. Additional 5 IPs and 5 TEPs were collected for RNA-seq, and 5 IPs and 4 TEPs were established the TOs. Exclusion criteria for all samples are: IP with EP histories, miscarriage, preeclampsia, and preterm delivery. EP with some risk factors associated with fallopian tubal abnormalities, *e.g.*, smoking, obvious tubal inflammatory adhesions, previous fallopian tubal diseases, and tubal surgery histories, were also excluded for avoiding complications. In each case, gestational days (GDs) was 40-64 days confirmed from the last menstrual period. Details about distribution of samples according to baseline characteristics are provided in Figure S1A and Table S1-3.

Samples collected by surgeries were processed immediately, including fixation in 10% formalin for histological and immunological analyses, immersed in RNAlater (Life Technologies) for following RNA-seq or real-time PCR analyses, and cultured in appropriate media for explant experiments. Samples for histological and immunological analyses were obtained and processed by the Department of Pathology, IPMCH following standard protocols briefed later.

**Hematoxylin-eosin staining (HE) of paraffin-embedded tissues**

The formalin-fixed, paraffin-embedded samples were cut into 5-μm slices using the microtome (Leica, RM2235), followed by staining with hematoxylin-eosin method by the Department of Pathology, IPMCH.

**Immunohistochemistry (IHC) of paraffin-embedded tissues**

Tissue blocks were cut into 5-μm slices as described before. Sections were deparaffinized, rehydrated and processed for antigen retrieval by a standard microwave heating technique in antigen retrieval solution (pH=8) (RecordBio, #RC016). Endogenous peroxidase activity was quenched by incubating the sections in 3% H_2_O_2_ for 25 min. The sections were incubated with 0.2% Triton X-100 and 3% bovine serum albumin (BSA) in PBS for 1 hour at room temperature and then incubated overnight with primary antibodies at 4 °C. Primary antibodies used for IHC include: monoclonal mouse anti-CD31 (PECAM-1) antibody (1:200, Abcam, #ab9498), monoclonal rabbit anti-CK7 antibody (1:200, ZSGB-BIO, #ZA-0573), and monoclonal rabbit anti-WNT2B antibody (1:100, Abcam, #ab211617). After overnight, samples were washed in PBS and incubated with the secondary antibody. Slides were visualized using diaminobenzidine tetrahydrochloride (RecordBio), and nuclei were counterstained with hematoxylin.

**Statistical analyses of villi and CD31-positive capillaries in cross sections**

In the CK7 of IHC sections, the 30 terminal villi of each section to be captured were selected at random, ensuring that it was analyzed blind by two independent observers. The measured parameters, including the mean cross area of villi, total cross area of villi and vascular numbers within the placental villi, were assessed by FIJI software (Wen et al., 2014).

**Immunofluorescence staining (IF)**

Primary antibodies used for IF include: monoclonal rabbit anti-podoplanin (D2-40) antibody (1:200, Bioss, #bs-21621R), monoclonal mouse anti-CD31 antibody (1:200, Abcam, #ab9498), monoclonal mouse anti-KI67 antibody (1:200, Cell Signaling Technology, #9449T), polyclonal rabbit anti-TEAD4 antibody (1:100, Cell Signaling Technology, #HPA056896), and polyclonal rabbit anti-CD138/Syndecan-1 (SDC-1) antibody (1:200, Abconal, #A4174). The sections were incubated overnight with primary antibodies at 4°C, followed by PBS washing and incubation with secondary antibodies. After washing with PBS, nuclei were stained with DAPI.

**Quantitative real-time PCR analysis**

Total RNA was extracted with RNA Isolator (Vazyme #R401-01). Equivalent amounts of RNA were reverse-transcribed using the HiScript II Q RT SuperMix (Vazyme #R223-01) according to manufacturer's instructions. cDNAs were then used as templates for each PCR reaction using ChamQ Universal SYBR qPCR Master Mix (Vazyme, #Q711-02) on QuantStudio 7 Flex Real-Time PCR system (Life Technologies). The expression level (relative quantity, RQ) of target mRNA was determined using the ^∆∆^Ct method with GAPDH/β-actin as the internal control with three replications. The sequences of gene-specific primers were listed in Table S5.

**Supplemental Tables**

| **Table S1** | Baseline characteristics of clinical samples in RNA-seq |
| --- | --- |
| **Table S2** | Baseline characteristics of clinical samples used to derive trophoblastic organoids |
| **Table S3** | Details of baseline characteristics of all clinical samples (XLS) |
| **Table S4** | The differentially expressed genes between IP and TEP (XLS) |
| **Table S5** | The primer sequences of RT-qPCR |
| **Table S6** | The material applied in the present study |

**Video S1-6. Video showing 3D renderings of the confocal stacks in Figure 1E and Figure S2C.**

| **Variables** | **IP group**  **(n=5)** | **TEP group**  **(n=5)** | ***P* value** |
| --- | --- | --- | --- |
| **Age^a^ (years)** | 32.00±3.35 | 34.80±2.46 | *P*=0.52 |
| **Gravidity^a^** | 1.80±0.58 | 2.20±0.37 | *P*=0.58 |
| **Parity^a^** | 0.60±0.24 | 0.60±0.24 | *P*=0.99 |
| **βhCG^a^ (IU/L)** | 16189.30±3718.00 | 6079.20±1093.00 | *P*=0.051 |
| **Gestational age^a^ (days)** | 45.20±1.59 | 44.20±1.77 | *P*=0.69 |
| **Fetus heart active (n, %)** | 4 (80.00) | 3 (60.00) | *P*=0.54 |
| **Pelvic adhesion (n, %)** | -- | 0 (0.00) | -- |

**Table S1. Baseline characteristics of clinical samples for RNA-seq.** βhCG: beta human chorionic gonadotropin.

^a^ Data are shown as the mean ± SEM. *P* values were calculated by Student's *t*-test.

| **Variables** | **IP group**  **(n=5)** | **TEP group**  **(n=4)** | ***P* value** |
| --- | --- | --- | --- |
| **Age^a^ (years)** | 31.40±3.84 | 29.50±1.19 | *P*=0.68 |
| **Gravidity^a^** | 1.60±0.40 | 2.25±0.48 | *P*=0.33 |
| **Parity^a^** | 0.40±0.24 | 0.50±0.29 | *P*=0.80 |
| **βhCG^a^ (IU/L)** | 19512.00±2921.00 | 5122.00±554.70 | *P*<0.01 |
| **Gestational age^a^ (days)** | 49.00±1.58 | 49.75±1.84 | *P*=0.77 |
| **Fetus heart active (n, %)** | 5 (100.00) | 3 (75.00) | *P*=0.29 |
| **Pelvic adhesion (n, %)** | -- | 0 (0.00) | -- |

**Table S2. Baseline characteristics of clinical samples for deriving trophoblastic organoids.** βhCG: beta human chorionic gonadotropin.

^a^ Data are shown as the mean ± SEM. *P* values were calculated by Student's *t*-test.

| **Species** | **Gene** | **Forward Primer** | **Reverse Primer** |
| --- | --- | --- | --- |
| Human | WNT1 | CGGCGTTTATCTTCGCTATC | TTCGATGGAACCTTCTGAGC |
|  | WNT2 | AAAGAAGATGGGAAGCGCCA | TTCATCAGGGCTCTGGCATC |
|  | WNT2B | CCTGTAGCCAGGGTGAACTG | CGGGCATCCTTAAGCCTCTT |
|  | WNT3 | CGCACGACTATCCTGGAC | GAGGCGCTGTCATACTTGTC |
|  | WNT3A | TGTTGGGCCACAGTATTCCT | GGGCATGATCTCCACGTAGT |
|  | WNT4 | CTAGCCCCGACTTCTGTGAG | TTGGACGTCTTGTTGCATGT |
|  | WNT5A | GCCCAGGTTGTAATTGAAGC | TGGCACAGTTTCTTCTGTCC |
|  | WNT5B | CGGGAGCGAGAGAAGAACT | CGTCTGCCATCTTATACACAGC |
|  | WNT6 | CAGCCCCTTGGTTATGGAC | AACTGGAACTGGCACTCTCG |
|  | WNT7A | AGAAGCAAGGCCAGTACCAC | GCCTCGTTGTTGTGCAAGTT |
|  | WNT7B | GCAGGAAGGTTCTAGAGG | GTTGTACTTCTCCTTCAGC |
|  | WNT8A | AGGCTGAGAAGTGCTACCAGA | CCATTGTTTGACCCATCACA |
|  | WNT8B | GAAGTACCACGCAGCACTCA | GAGATGGAGCGAAAGGTGTC |
|  | WNT9A | GGTGTGAAGGTGATCAAG | TGCCGTCTCATACTTGTG |
|  | WNT9B | AGAGGAAACAAGGACCTGCG | GCACTTACACGTGGTCCTGA |
|  | WNT10A | CATCTTCAGCAGAGGTTTC | CAGTGCATCCAGTTGTAAG |
|  | WNT10B | GATACCCACAACCGCAATTC | GGGTCTCGCTCACAGAAGTC |
|  | WNT11 | GGCCAAGTTTTCCGATGCTC | CACCCCATGGCACTTACACT |
|  | WNT16 | GAAACTGGATGTGGTTGG | TCATGCAGTTCCATCTCTC |
|  | VEGF | TGCAGATTATGCGGATCAAACC | TGCATTCACATTTGTTGTGCTGTAG |
|  | β-Actin | CATGTACGTTGCTATCCAGGC | CTCCTTAATGTCACGCACGAT |
|  | GAPDH | GTCTCCTCTGACTTCAACAGCG | ACCACCCTGTTGCTGTAGCCAA |

**Table S5. The primer sequences of RT-qPCR.**

**Supplementary Figures**

| **Figure S1** | Statistics and comparison of IP, AEP and TEP villi and fetuses. |
| --- | --- |
| **Figure S2** | Placental intravillous vascularization in IP, AEP and REP. |
| **Figure S3** | Transcriptome analyses of signaling pathways and developmental states in IP and TEP conditions. |
| **Figure S4** | Secretory factors from IP placental villous explants promote angiogenesis in a WNT-dependent manner. |
| **Figure S5** | VEGF-mediated angiogenesis is dependent on WNT signaling but less dependent on WNT ligand-receptor interaction. |
| **Figure S6** | Trophoblast organoids (TOs) derived from IP and TEP placental villi exhibit similar WNT2B expression level. |

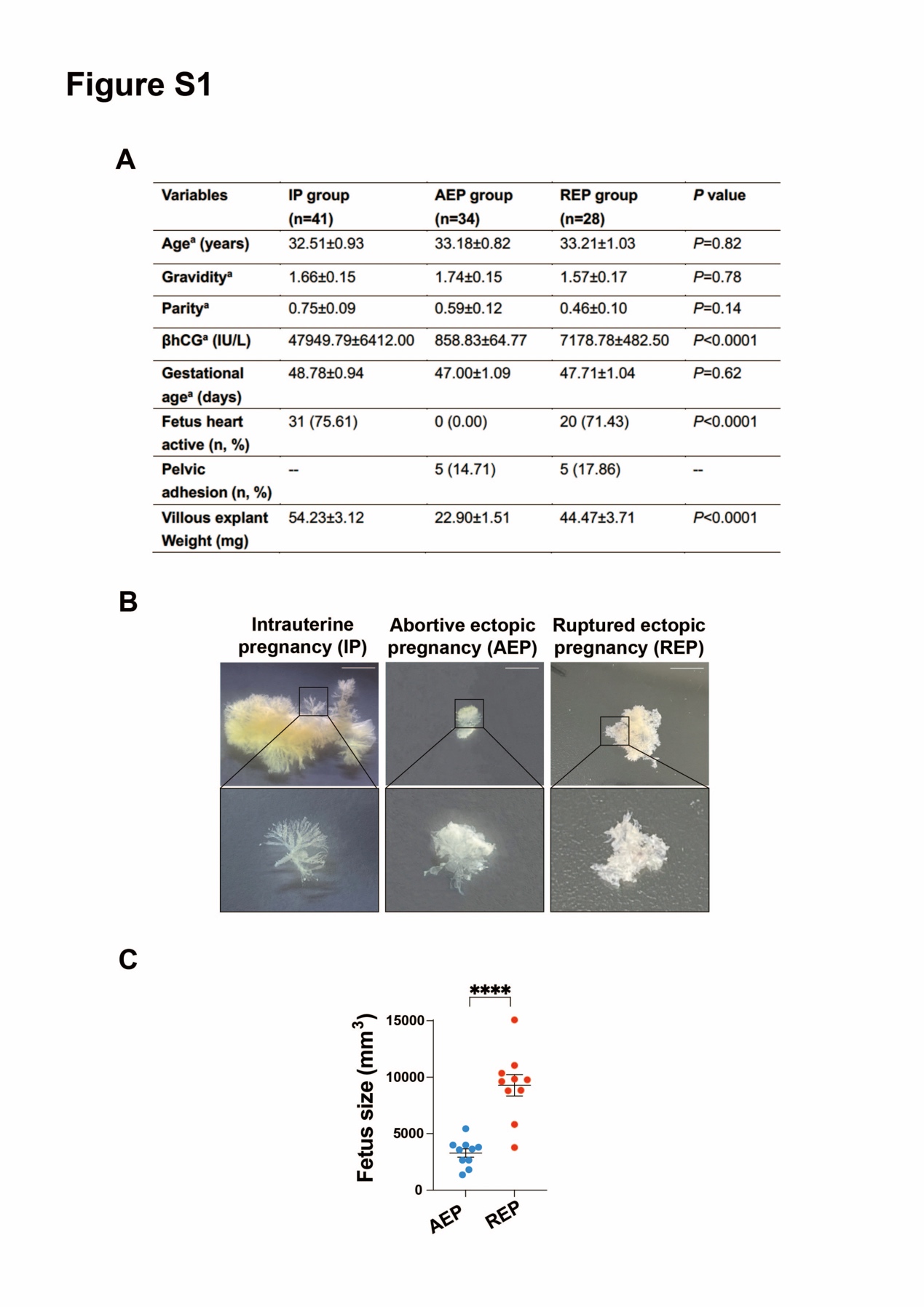

**Figure S1. Statistics and comparison of IP, AEP and TEP villi and fetuses.** (A) Baseline characteristics of clinical samples in three groups. βhCG: beta human chorionic gonadotropin. Data are shown as the mean ± SEM. *P* values were calculated by one-way ANOVA, Tukey’s test. (B) The morphologies of placental villi from IP, AEP and REP. Compared with AEP and REP, the volume of placental villi from IP is much larger, with obvious dendritic structures. In REP, volume of placental villi is 2-3 times larger than that in AEP. Scale bars, 1cm. (C) Comparison of sizes of AEP and REP fetuses as measured by ultrasound imaging.

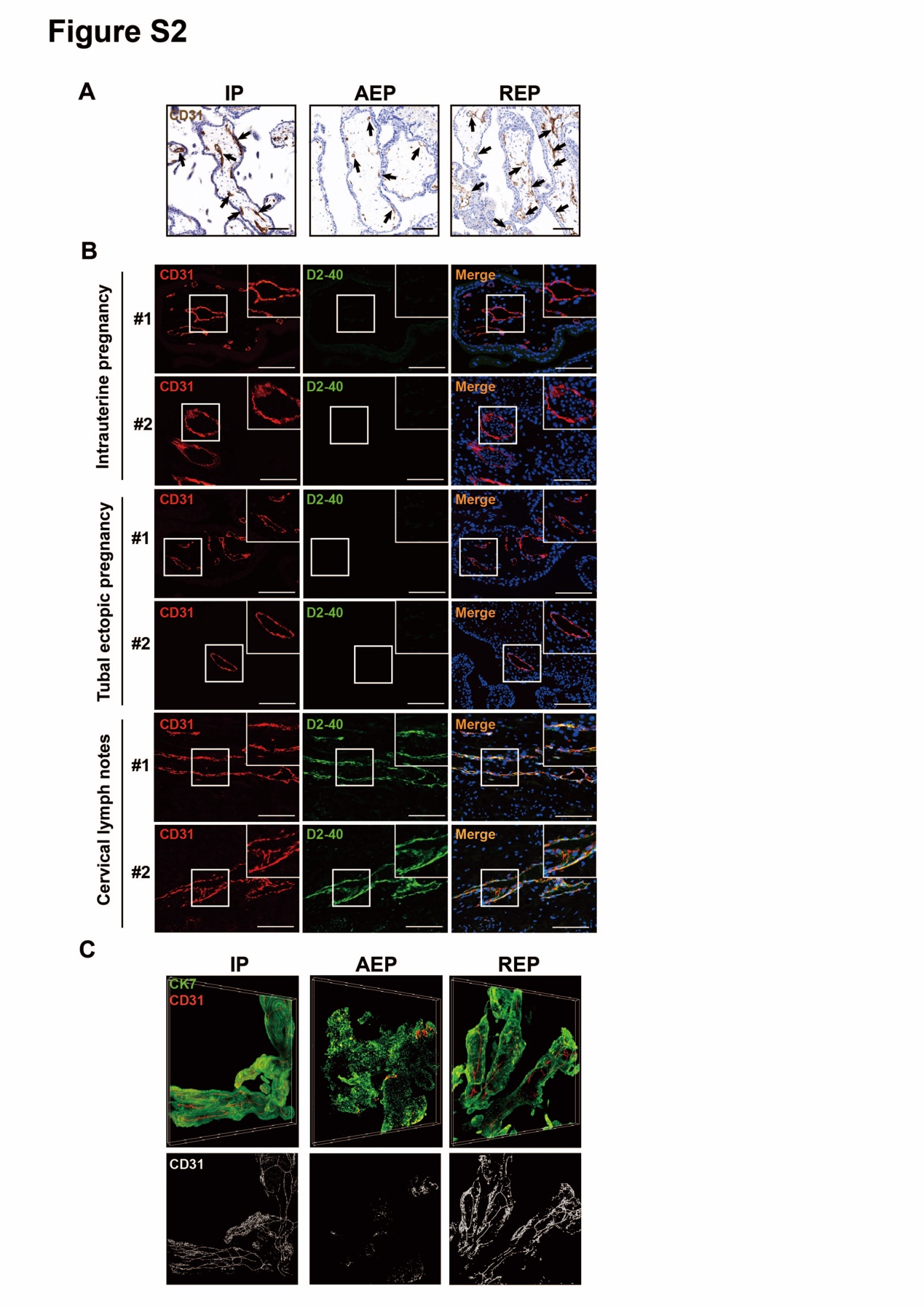

**Figure S2. Placental intravillous vascularization in IP, AEP and REP.** (A) Immunohistochemical staining of CD31 in longitudinal sections of IP, AEP, and REP placental villi. Relevant to Figure 1D to show additional clinical samples. Scale bars, 100 μm. (B) Absence of lymphatic endothelium marker in early placental villi. The cervical lymphatic tissues are positive for CD31 (red) and D2-40 (green). Whether it is IP villi or EP villi, both are shown CD31 (+) and D2-40 (-). Nuclei are labelled with DAPI (blue). Scale bars, 100 μm. (C) IP and REP placental villi exhibit higher vascularization level in cleared tissues. IP, AEP, and REP placental villi were cleared and immunostained for CK7 (green, trophoblasts) and CD31 (red, endothelial cells). Stereoscopic views for each type of pregnancy are shown. Lower, skeletonization of CD31 staining to visualize vascular systems of IP, AEP and REP placental villi. Relevant to Figure 1E to show additional clinical samples.

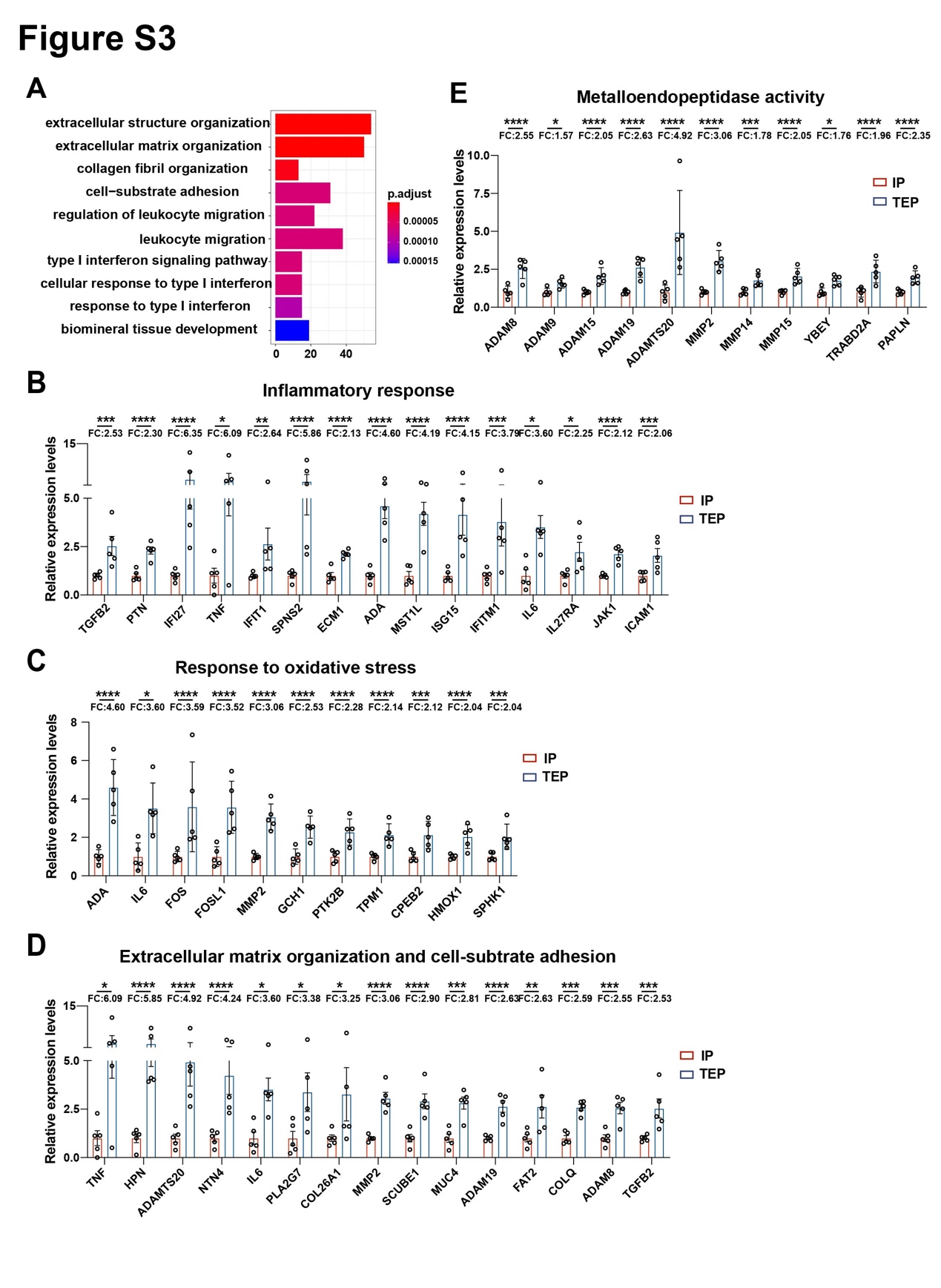

**Figure S3. Transcriptome analyses of signaling pathways and developmental states in IP and TEP conditions.** Gene Ontology (GO) analyses reveals the pathway-specific enrichment of genes up-regulated in TEP, including metalloendopeptidase activity, oxidative stress the response, inflammatory responds, and extracellular matrix organization and cell-substrate adhesion. Log-transformed, normalized expression levels for selected typic markers present differentially up-regulated genes between IP (n=5) and TEP (n=5). Data represents mean ± SEM. *P* values were calculated by student's *t*-test or Welch's *t*-test.

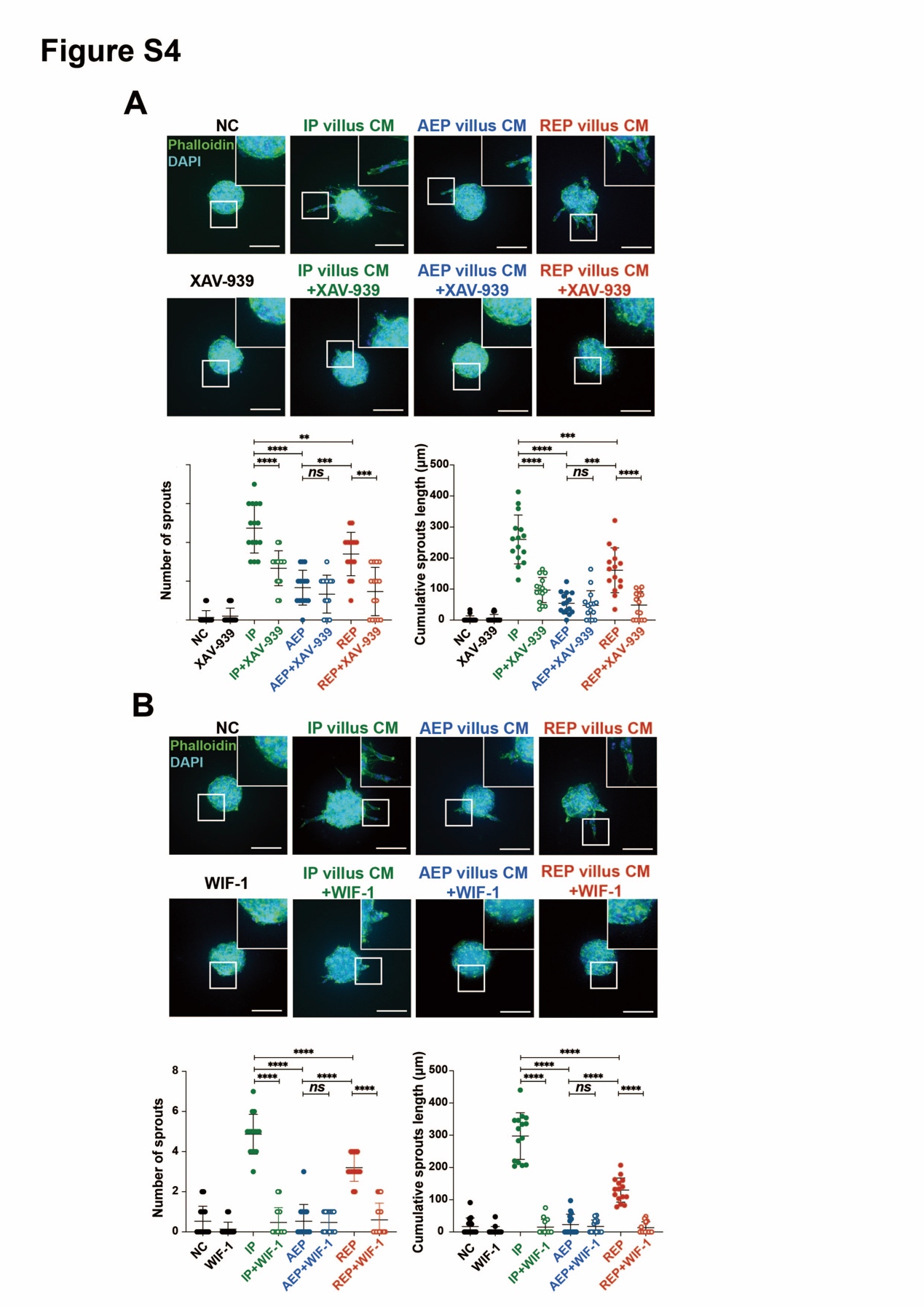

**Figure S4. Secretory factors from IP placental villous explants promote angiogenesis in a WNT-dependent manner.** In sprouting angiogenesis assay, conditioned media from three types of pregnancies were co-treated with varies inhibitors antagonizing different components of canonical WNT signaling pathway, such as XAV-939 (4 μM) (A) or WIF-1 (2.5 μg/ml) (B). The number of sprouts per spheroid and the cumulative sprout length indicate that WNT inhibitors greatly abolish pro-angiogenic effects of conditioned media from IP and REP villous explants. Data represents mean ± SD. Scale bars, 100 μm. **P* < .05, ***P* < .01, ****P* < .001, *****P* < .0001. *P* values were calculated by ANOVA, Tukey’s test and student's *t*-test or Welch's *t*-test. NC: negative control.

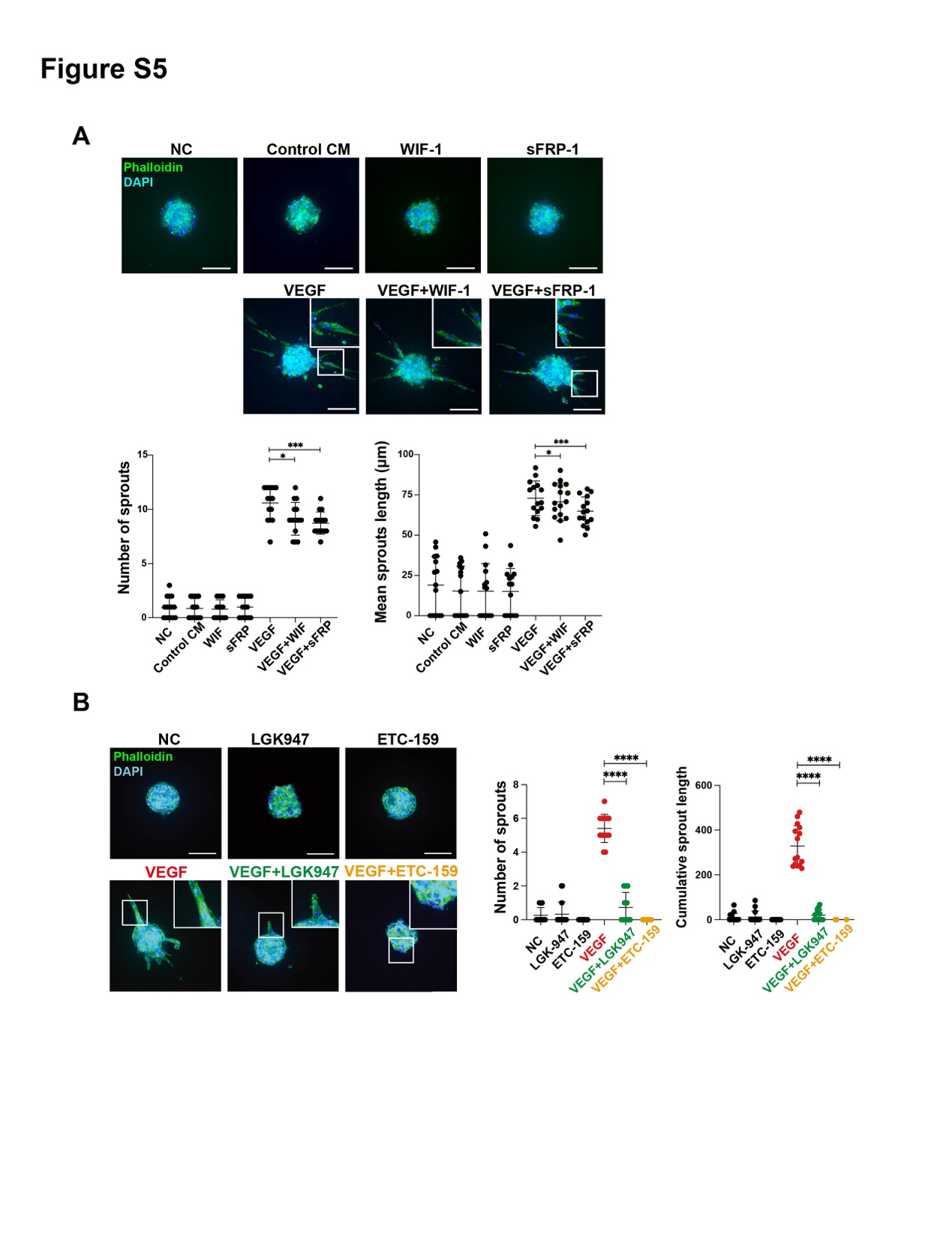

**Figure S5. VEGF-mediated angiogenesis is dependent on WNT signaling but less dependent on WNT ligand-receptor interaction.** (A) Capillary-like sprouting angiogenesis was quantified after 24 h of stimulation with VEGF (250 ng/ml) the presence or absence of sFRP (5 μg/ml) or WIF-1 (2.5 μg/ml). Lower, the statistical data of number of sprouts per spheroid and cumulative sprout length indicates that sFRP-1 and WIF-1 only slightly blocked the pro-angiogenic effects of VEGF. (B) PORCN inhibitors LGK947 (10 ng/ml) and ETC-159 (100 ng/ml) largely abolished VEGF-induced sprout formation of HUVECs. Data represents mean ± SD. Scale bars, 100 μm. **P* < .05; ****P* < .001. *P* values were calculated by ANOVA, Tukey’s test, student's *t*-test or Welch's *t*-test. NC, negative control.

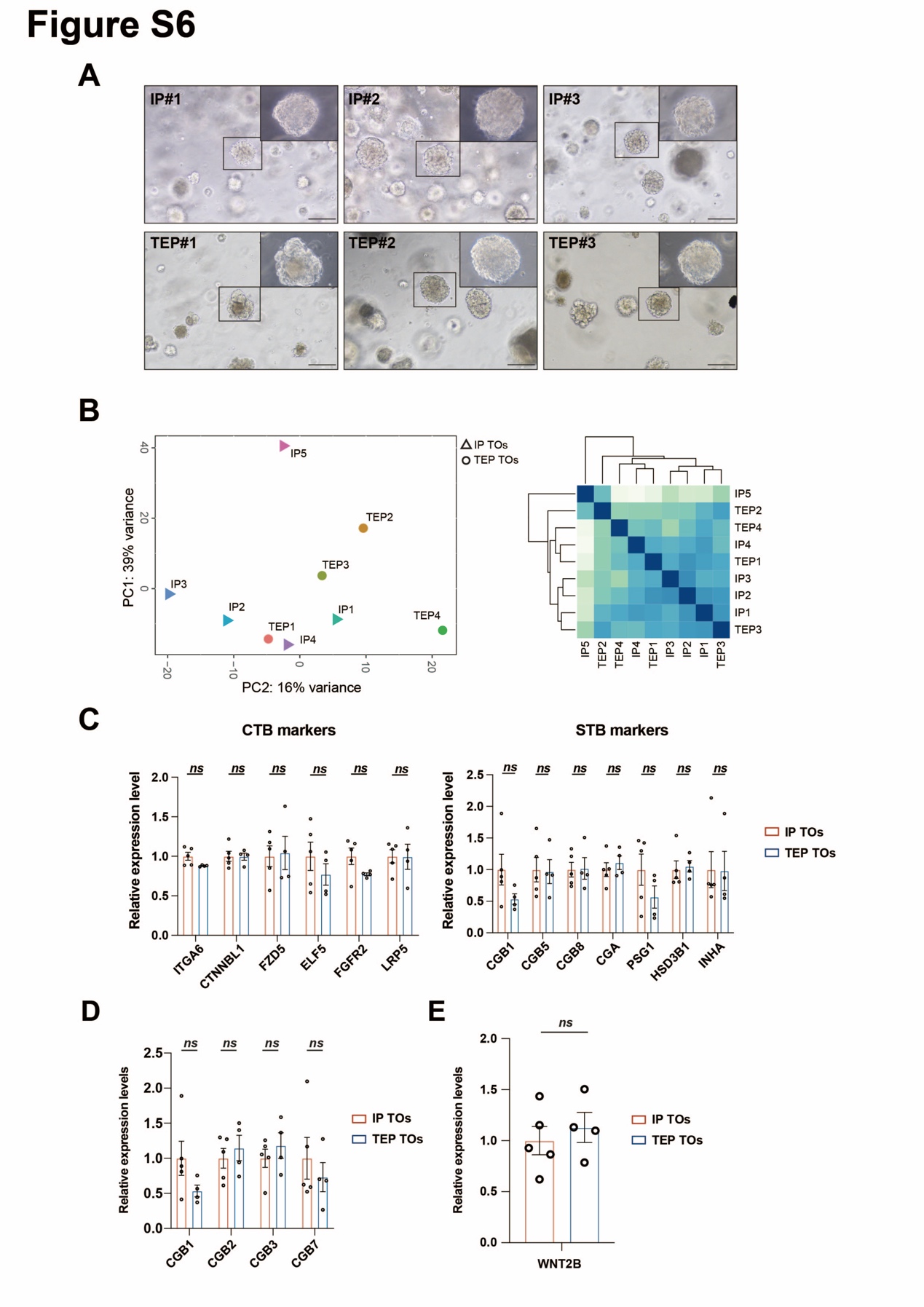

**Figure S6. Trophoblast organoids (TOs) derived from IP and TEP placental villi exhibit similar WNT2B expression level.** (A) Phase-contrast images of trophoblastic organoids in the IP (n=3) and TEP (n=3). Scale bars, 100 μm. (B-C) Unsupervised principal component analysis (PCA) and Pearson analysis suggest no significant clustering of TEP (n=4, circle) and IP TOs (n=5, triangle) in RNA-seq analysis. (D-E) No significant differences in expression of CTB and STB markers, hCG, and WNT2B between IP and TEP TOs. Log-transformed, normalized expression levels of selected genes are retrieved from RNA-seq analysis. Data represents mean ± SEM. *P* values were calculated by student's *t*-test or Welch's t-test.
